# Characterization of a sex-determining region and its genomic context via statistical estimates of haplotype frequencies in daughters and sons sequenced in pools

**DOI:** 10.1101/2020.08.06.240416

**Authors:** Richard Cordaux, Mohamed Amine Chebbi, Isabelle Giraud, David Pleydell, Jean Peccoud

## Abstract

Sex chromosomes are generally derived from a pair of autosomes that have acquired a locus controlling sex. Sex chromosomes usually evolve reduced recombination around this locus and undergo a long process of molecular divergence. Although sex chromosomes have been intensively studied in several model taxa, the actual loci controlling sex are difficult to identify in highly diverged sex chromosomes, hence they are known in relatively few species. Taxa with evolutionarily young sex chromosomes can help fill this gap in knowledge. Here we aimed at pinpointing the sex-determining region (SDR) of *Armadillidium vulgare*, a terrestrial isopod with female heterogamety (ZW females and ZZ males) and which presumably presents evolutionarily young sex chromosomes. To locate the SDR, we assessed SNP allele frequencies in F1 daughters and sons sequenced in pools (pool-seq) in several families. We developed a Bayesian method that uses the SNP genotypes of individually sequenced parents and poolseq data from F1 siblings to estimate the genetic distance between a given genomic region (contig) and the SDR. This allowed us to assign more than 43 Megabases of contigs to sex chromosomes. By taking advantage of the several F1 families, we delineated a very short genomic region (~65 kilobases) that did not show evidence for recombination with the SDR. In this region, the comparison of sequencing depths between sexes outlined female-specific genes that may be involved in sex determination. Overall, our results provide strong evidence for an extremely low divergence of sex chromosomes in *A. vulgare*.

## Introduction

The existence of males and females (gonochorism) constitutes a phenotypic variation found in many taxa and which has a profound impact on their evolution. Despite gonochorism being common and ancient, the mechanisms initiating the developmental cascade toward the male or female phenotype appear to be highly variable (BACHTROG *et al.* 2014; BEUKEBOOM AND PERRIN 2014). In some species, sex is solely or partially determined by environmental factors such as temperature (MERCHANT-LARIOS AND DIAZ-HERNANDEZ 2013) and social interactions (BRANTE *et al.* 2016) while in many others, sex is entirely determined by genotype (review in BACHTROG *et al.* (2014); BEUKEBOOM AND PERRIN (2014)).

When the sex of individuals is genetically determined, it is generally under the control of sex chromosomes. In the strict sense, sex chromosomes are chromosomes that carry a gene (in the mendelian sense) whose genotype determines the sex of its carrier. Sex chromosomes are not required to be a pair of chromosomes that look different from each other under the microscope, a property that is referred to as heteromorphy. In fact, sex chromosomes should initially be homomorphic as they are usually derived from a pair of autosomes that have acquired a sex determining locus (WRIGHT *et al.* 2016; FURMAN *et al.* 2020). As opposed to homologous autosomes however, sex chromosomes generally diverge under balancing selection and the influence of alleles with sex-antagonistic effects (BACHTROG *et al.* 2014; WRIGHT *et al.* 2016). This evolution generally leads to reduced crossing over rates around the master sex-determining gene (BERGERO AND CHARLESWORTH 2009). In effect, the non-recombining chromosomal region that associates with the sex phenotype (hereafter referred to as the sex-determining region or SDR), tends to increase in size. The reduction of recombination leads to the divergence of the two chromosomes, due to inability to efficiently purge deleterious mutations (BERGERO AND CHARLESWORTH 2009; BACHTROG 2013). Through this incessant divergence process, sex chromosomes may become visually recognizable in a karyotype, which enabled the identification of different genetic sex-determining systems (reviewed in BACHTROG *et al.* (2014)). The most notorious ones are the XY system where males are heterogametic (i.e., they are heterozygous at the sex-determining gene) (as in therian mammals and *Drosophila*), and the ZW system where females are heterogametic (as in birds and lepidopterans). The ease to identify a pair of heteromorphic sex chromosomes is however balanced by the difficulty to locate the master sex-determining gene, as this gene is only part of a large non-recombining SDR that may occupy most of the chromosome length. This difficulty may explain why the identified sex-determining genes are still few in comparison to the diversity of taxa with chromosomal sex determination, as most model organisms (mammals, birds, fruit flies, etc.) harbor highly heteromorphic sex chromosomes. In taxa where sex chromosomes appear to undergo rapid turnover, such as teleost fishes (MANK AND AVISE 2009), sex chromosomes are evolutionarily young, hence SDRs likely to be short. Short SDRs have helped to pinpoint several sex-determining genes (e.g. KAMIYA *et al.* (2012); AKAGI *et al.* (2014)), some of which vary among related species within the same genus (MATSUDA *et al.* 2002; NANDA *et al.* 2002). These taxa therefore emerge as useful models to study the appearance and early evolution of sex chromosomes (CHARLESWORTH *et al.* 2005) and to better characterize the nature and diversity of sex-determining genes, hence the reaction cascades leading to the development of sex phenotypes.

Terrestrial isopods, also known as woodlice or pillbugs, are good biological models for this endeavor. These crustaceans indeed appear to undergo a dynamic evolution in their sex chromosomes (JUCHAULT AND RIGAUD 1995), as evidenced by multiple transitions between XY systems and ZW systems (BECKING *et al.* 2017). These recurrent transitions imply that sex chromosomes of several isopod species may be evolutionarily young. Among these species, the common pillbug *Armadillidium vulgare* has been the most studied with respect to sex determination (CORDAUX *et al.* 2011). Although sex chromosomes are visually undistinguishable among the 27 chromosome pairs composing the *A. vulgare* genome (ARTAULT 1977), crossing experiments using genetic females masculinized via hormones (JUCHAULT AND LEGRAND 1972) have demonstrated that this species presents ZW sex chromosomes. Homomorphy of the *A. vulgare* sex chromosomes is also consistent with the reported viability and fertility of WW individuals generated through these crossing experiments. The molecular similarity between the Z and W chromosomes of *A. vulgare* has more precisely been evaluated through the assembly and analysis of its ~1.72 Gbp genome (CHEBBI *et al.* 2019). More specifically, comparison of sequenced depths from data obtained from males and females allowed locating W-specific sequences (those with no similarity found in ZZ males) in 27 contigs representing less than 1 Mbp (CHEBBI *et al.* 2019). However, the *A. vulgare* SDR might not correspond to a region of absence of similarity between Z and W chromosomes. In some species, sex has been shown to be controlled by a single nucleotide polymorphism (SNP) (KAMIYA *et al.* 2012), a hypothesis that cannot be excluded in *A. vulgare*. In effect, very little is known about the *A. vulgare* sex chromosomes beyond the 27 contigs containing W-specific sequences.

In this situation, methods based on SNPs can be useful as they can locate SDRs, and genetically linked loci, in sex chromosomes presenting very low molecular divergence. Researchers would locate loci for which individuals of a given sex all present expected genotypes (e.g., loci presenting SNPs that are heterozygous in all ZW daughters and homozygous in all ZZ sons). However, when no information on the location of the SDR is available (e.g., in the absence of genetic map) and if no candidate genes for sex determination are suspected, the whole genome must be analyzed, hence re-sequenced or scanned. Such endeavor can be very costly if whole-genome sequencing is undertaken on many individuals, especially considering the sample size required for reliable statistical inference. These considerations motivated the use of techniques permitting partial genome sequencing (PALMER *et al.* 2019), such as Restriction-site Associated DNA markers (RAD sequencing) (BAIRD *et al.* 2008), which reduce sequencing costs at the expense of an increase in DNA library preparation costs. An alternative strategy consists of pooling the DNA from multiple individuals prior to sequencing (FUTSCHIK AND SCHLÖTTERER 2010), a method referred to as “pool-seq”. Pool-seq drastically cut DNA library preparation costs, and generally sequencing costs as well given that the sequencing effort per individual is generally lower than in the case of individual whole genome sequencing. Pool-seq substitutes the obtention of individual genotypes with estimates of allele frequencies within pools. The main drawback is that allele frequencies among the pooled individuals are inferred from allele frequencies among sequenced fragments (“reads”) covering a SNP, adding some degree of uncertainty that diminishes with increased sequencing effort (GAUTIER *et al.* 2013). As each SNP is analyzed separately, somewhat arbitrary thresholds on estimated frequencies among reads and on sequencing depths are used to determine whether individual SNPs are associated to the sex phenotype (e.g., PAN *et al.* (2019)). Also, the absence of individual genotypes prevents the construction of a genetic map, limiting knowledge about the genomic context of loci of interest.

Here, we developed a statistical approach that tackles these limitations, based on the individual genotyping of parents and pool-seq of progeny from several crosses. This method allowed us to identify a short genomic region that likely contains the SDR of *A. vulgare*. Beyond the SDR, our approach allowed us to assign more than 43 Mb of contigs to sex chromosomes, even though these chromosomes appeared no different from autosomes with respect to molecular divergence and showed uniform recombination rates.

## Materials and methods

### General approach

We developed a pool-seq based approach to locate a ZW-type SDR by considering that a W-linked allele (where “linked” here denotes the absence of recombination with the SDR) should have frequencies of 0.5 in females and 0 in males. This W-linked allele has a Z-linked counterpart with frequencies of 0.5 in females and 1 in males. We estimate allele frequencies in daughters and sons from the F1 progeny of two parents through whole-genome sequencing of pools of individuals of the same sex, followed by read mapping on a genome assembly (Figure 1). Our analysis relies on biallelic SNPs for which the mother is heterozygous and the father homozygous, hereafter called “informative SNPs”. Allele frequencies at SNPs that are heterozygous in the father depend on random sampling of paternal gametes, which do not determine the sex of the progeny.

**Figure 1.**
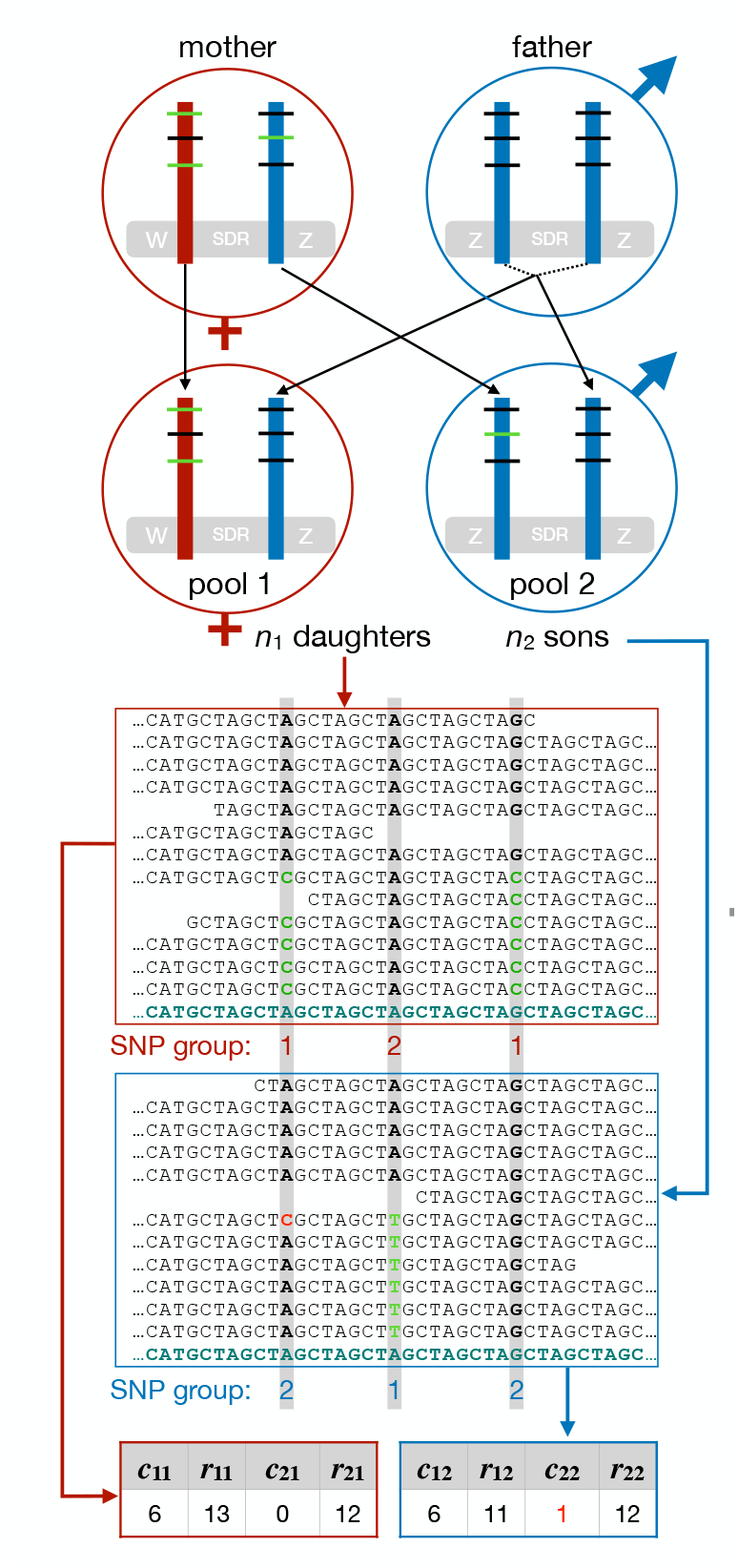
Locating the ZW sex determining region (SDR) via a cross where the genomes of several F1 siblings per sex are sequenced in pools and parental genomes sequenced individually. Red/blue rods represent regions of W/Z chromosomes possessing informative SNPs shown as horizontal segments. For each SNP, the maternal allele (as defined in the methods section) appears in green and the paternal allele in black. In this example, no crossing over occurred between the SNPs and the SDR. The mapping of reads obtained from the pools shows sequences (reads) aligned on the reference genome (bottom sequence), with three informative SNPs outlined. A sequencing error is shown in red. Tables at the bottom show the values of the variables used to estimate the frequency of the focal haplotype (the W-carried haplotype for daughters and the Z-carried haplotype for sons), based on the mapped reads. See section “Statistical estimation of haplotype frequencies” for the definition of these variables.

Rather than estimating allele frequencies in F1s for each individual SNP, we estimate the frequency of haplotypes combining the alleles of all informative SNPs of a given locus (genomic contig). This allows much greater precision in our estimates since SNP alleles of the same haplotype have the same frequency in a pool, assuming the absence of crossing over within the contig. We aim at identifying the haplotypes carried by maternal chromosomes and whose frequencies in an F1 pool depend on the pool’s sex.

Maternal haplotypes are defined as follows. We first consider that the allele that is specific to the mother at an informative SNP, hereafter called “maternal allele”, can be carried either by the W or Z chromosome (Figure 1). We define as “type-1” a SNP whose maternal allele is carried by the maternal W chromosome, and as “type-2” a SNP whose maternal allele is carried by the maternal Z. Hence, the W-linked haplotype carries maternal alleles of type-1 SNPs and paternal alleles of type-2 SNPs. To reconstruct this haplotype, we use the fact that W-linked maternal alleles are predominantly inherited by daughters, and that Z-linked maternal alleles are predominantly inherited by sons (Figure 1). Hence, we classify a SNP as type 1 whenever the maternal allele is more frequent in daughters than in sons, and as type 2 in the opposite situation. If the maternal allele is equifrequent in both sexes, we attribute types at random.

To enable the same mathematical development for both sexes, we introduce the notion of SNP “group”. SNP group is the same as SNP type in daughters, while we assign every type-1 SNP to group 2 and every type-2 SNP to group 1 in sons. Each SNP is thus assigned to different groups in different sexes of the progeny. This allows us to define the “focal haplotype” as the one that combines the maternal alleles of group-1 SNPs and the paternal alleles of group-2 SNPs, regardless of the sex. The focal haplotype is thus the W-linked haplotype in daughters and the Z-linked maternal haplotype in sons. In the absence of recombination with the SDR, the focal haplotype must be inherited by all the descendants of a given sex, hence its frequency must be 0.5 in a pool (the remainder represents paternal DNA).

### Statistical estimation of haplotype frequencies

In a given pool *i* of *n*_*i*_ individuals born from *n*_*i*_ oocytes, we denote as *f* the unknown frequency of the focal haplotype at a locus, with potential values in the set ***F***_***i***_ = {0…*n*_*i*_ / (2*n*_*i*_)}. To put *f* in perspective, the proportion of oocytes that underwent recombination between the contig and the SDR, which is the distance to the SDR in Morgans, is 1 − 2*f*. We then let *c*_1*i*_ denote the number of sequenced DNA fragments (e.g., read pairs) from pool *i* carrying the maternal alleles of all group-1 SNPs of the locus. We let *r*_1*i*_ represent the number of fragments from pool *i* covering group-1 SNPs (both alleles combined). We finally let *c*_2*i*_ and *r*_2*i*_ denote equivalent variables for group-2 SNPs. To shorten the notation, we refer to the set of variables {*c*_1*i*_, *r*_1*i*_, *c*_2*i*_, *r*_2*i*_} as ***cr***_***i***_.

These specifications allow expressing the posterior probability of the focal haplotype frequency given the observed data, via Bayes’ theorem:

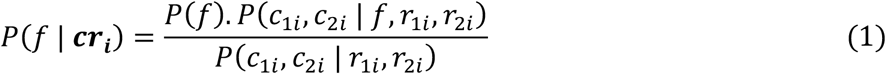

We use a uniform prior for *f*, hence *P*(*f*) = 1/(*n*_*i*_ + 1) ∀ *f* ∈ ***F***_***i***_. Although a prior based on *Binomial*(*n*_*i*_, 0.5) would more accurately represent the inheritance of maternal haplotypes for most contigs, such prior would not be suited for contigs at less than 50 centiMorgans (cM) to the SDR, the frequency of which is unknown. In particular, a binomial prior assumes that *f* is quite unlikely to equal 0.5 (with a probability of 0.5^*ni*^), increasing the risk of false negatives when it comes to the selection of contigs linked to the SDR. We would rather include false positives in our selection of candidates, as this selection is only a first step in the search for the sex-determining gene(s). To specify *P*(*c*_1*i*_, *c*_2*i*_ | *f, r*_1*i*_, *r*_2*i*_), we assume the sequencing of the different SNP groups of a contig to be independent. Although this is not true if SNPs of different groups are covered by the same read pairs, we consider our approximation to be acceptable and stress the difficulty of taking this dependence into account. Hence,

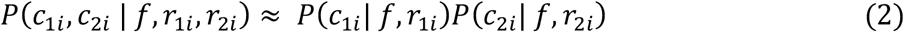

We then consider *c*_1*i*_ as the realization of *Binomial*(*r*_1*i*_, *f*), i.e., that the *c*_1*i*_ read pairs carrying the maternal allele of group-1 SNPs result from *r*_1*i*_ independent draws among DNA molecules containing the focal haplotype, which has frequency *f* in the pool. *c*_2*i*_ is the realization of *Binomial*(*r*_2*i*_, 0.5 − *f*) as the maternal alleles group-2 SNPs are carried by the non-focal maternal haplotype, which constitutes the rest of the maternally inherited chromosomes, themselves constituting half of the chromosomes in the F1. For a contig linked to the SDR, *c*_2*i*_ should be zero (Figure 1). However, a nucleotide indicating the maternal allele in a read may result from an “error”. Errors combine mutations between parents and offspring, *in vitro* mutations and sequencing or mapping errors. An error causing *c*_2*i*_ to be positive would nullify the posterior probability that *f* = 0.5, hence that no crossing over occurred between the locus and the SDR. To avoid this, we introduce a constant ε to represent the rate of error leading to a maternal allele appearing in a read. The probability that a read carries the maternal allele of a group-2 SNP becomes (1 − ε)(1/2 − *f*) + ε = (1 + ε)/2 + ε*f* −*f*.

We do not apply this correction to group-1 SNPs. A group-1 SNP of a contig linked to the SDR should present both alleles at the same frequency, such that errors leading to the detection of maternal alleles are compensated by error leading to the detection of non-maternal alleles.

From the above considerations,

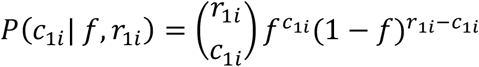

and

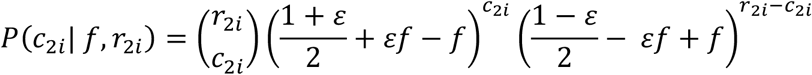

The marginal probability of the maternal allele read counts, *P*(*c*_1*i*_, *c*_2*i*_ | *r*_1*i*_, *r*_2*i*_), integrates over all the values that *f* can take, hence

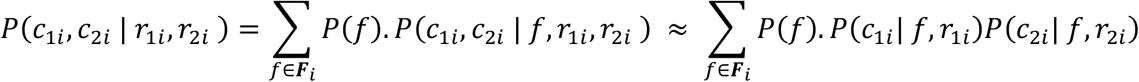

Replacing terms in equation 1 yields the following, after cancellation of terms present in the numerator and denominator:

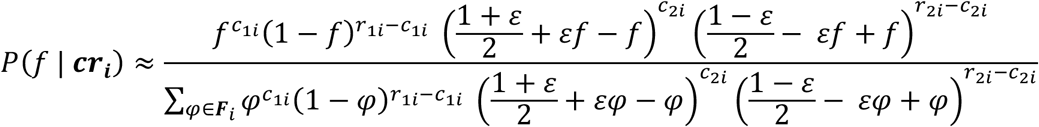

These posterior probabilities were used to estimate the expected number of recombination events, denoted here as *n*_rec_, that occurred between a given contig and the SDR in the oocytes of several pools. This was done by calculating the sum of the expected number of recombination events over pools as follows:

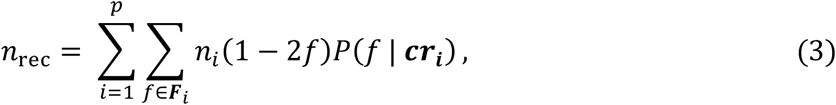

where *p* is the number of pools. This formula accounts for variable levels of uncertainty in the true value of *f* among pools. Note that while the true number of recombination events is a discrete random variable, its expected value *n*_rec_ is a weighted average and is therefore a continuous variable.

Importantly, the accuracy of *n*_rec_ depends on the ability to reconstruct the maternal haplotypes, via inference of SNP types (see previous section). This ability decreases with the genetic distance to the SDR, which leads to underestimating of the number of recombination events (Supplementary text, Figure S1). In our experiment, this bias becomes noticeable at about 8 recombination events, corresponding to ~20 cM to the SDR, but it is minimal on contigs locating the closest to the SDR (Figure S1).

### Crosses, sequencing and mapping

We applied our approach to three *A. vulgare* lines: WXa, ZM and BF (Table 1), which have been shown to have homologous sex chromosomes (CHEBBI *et al.* 2019). For each line, a single virgin female was crossed with a single male until it showed evidence for gravidity and then isolated to lay progeny. DNA was extracted from gonads, heads and legs of *n* = 10 descendants of each sex with the Qiagen blood and tissue kit, according to the protocol for animal tissues (3h of incubation in proteinase K at 56°C and 30 min of RNase treatment at 37°C). Absence of *Wolbachia* endosymbionts and the *f* element in all samples was confirmed by PCR, as described previously (LECLERCQ *et al.* 2016). DNA concentration was estimated for each sample by Qubit fluorometric quantification, to enable pooling DNA samples in equimolar proportions. DNA samples from 10 individuals per sex constituted a pool containing 7 μg of DNA. Each pool was sequenced on an Illumina HiSeq2500 platform (125-bp paired ends) by Beckman Coulter Genomics. We aimed at a sequencing depth of 30X per pool to ensure that most parts of the 20 chromosome doses (i.e. from 10 diploid individuals) in the pools were sequenced. The DNA of individual parents from lines WXa and ZM was extracted as described above and sequenced on an Illumina HiSeqX platform (150-bp paired ends) by Génome Québec. To enable reliable genotyping, we targeted an average sequencing depth of 30X per parent. However, technical reasons unrelated to our approach prevented the sequencing of parents from line BF.

**Table 1.** Characteristics of the three *A. vulgare* lines used in this study.

|  |  |  |  |
| --- | --- | --- | --- |
| Line/family | WXa/1591 | ZM/544 | BF/2875 |
| Source location | Helsingør, Denmark | Heraklion, Greece | Nice, France |
| Collection date | 1982 | 1989 | 1967 |
| GenBank accession numbers: |  |  |  |
| - Father (ZZ) | Xxx | Xxx | Not sequenced |
| - Mother (ZW) | Xxx | Xxx | Not sequenced |
| - Pool of 10 sons (ZZ) | Xxx | Xxx | SRR8238987 |
| - Pool of 10 daughters (ZW) | xxx | xxx | SRR8238986 |

Xxx : GenBank accession numbers will be provided at revision stage.

Sequencing reads were trimmed from low-quality parts using trimmomatic version 0.33 (BOLGER *et al.* 2014). For each F1 pool and parent, trimmed reads were aligned on the female reference *A. vulgare* genome (CHEBBI *et al.* 2019) using bwa_mem (LI 2013) with default settings. In the resulting alignment (bam) file, reads sequenced from the same original DNA fragments (PCR or optical duplicates) were flagged by picardtools MarkDuplicates version 2.12.0 (http://broadinstitute.github.io/picard/, last accessed June 9 2020). Reads containing indels were realigned on the reference genome using GATK’s IndelRealigner. To establish the whole-genome genotype of each parent, we followed the GATK best practices (VAN DER AUWERA *et al.* 2013) as described in CHEBBI *et al.* (2019). This involved recalibrating base quality scores of mapped reads to reduce the risk of considering sequencing errors as variants, followed by SNP genotype calling with HaplotypeCaller. Genotyping was performed independently on each parent, recording all positions in a gvcf file. The four gvcf files were merged using GenotypeGvcf in a single vcf file, discarding putative SNPs not passing quality check. We used this file to select informative SNPs, excluding positions with indels, lack of sequence data in any individual or with more than two alleles.

### Estimation of haplotype frequencies within F1 pools

We used a custom R script (R DEVELOPMENT CORE TEAM 2020) that scans each F1 bam file via samtools version 1.10 (LI *et al.* 2009) to retrieve the base carried by each read at informative SNPs, associated with a unique read-pair identifier. Read pairs marked as duplicates were ignored as well as secondary alignments and those with mapping quality score <20. For each pool and SNP, we counted reads carrying the maternal and paternal alleles. Reads carrying other alleles were ignored. At each informative SNP, the frequency of the maternal allele in the pool was estimated by the proportion of reads carrying this allele. By comparing this frequency between sexes of the same family, we inferred SNP type and group (1 or 2) using the principles described earlier. For each contig in each pool *i*, we established the read count data (variables of the ***cr***_***i***_ set) by counting read pairs according to the definition for these variables. We then computed *P*(*f* | ***cr***_***i***_) for every possible value of *f*. We set the error rate ε at 0.01, which is higher than the typical sequencing error rate of the Illumina technology. Using results from the four pools of the WXa and ZM lines (for which we sequenced the parents), we estimated the number of recombination events between the contig and the SDR, *n*_rec_. For these computations, we excluded any SNP that failed to pass the following criteria, which we applied independently for both families. Genotyping quality in the mother (determined by GATK’s haplotype caller) had to be higher than 10 and that of the father higher than 40 (the presence of the rarer maternal allele in the F1 allowed us to be less restrictive on the mother’s genotype quality, while we wanted to ensure that the father was not homozygous). In addition, at least one read had to carry the maternal or paternal allele in each F1 pool, the total number of reads carrying either allele in both F1 pools combined had to not exceed its 95% quantile (excessive sequencing depth may reflect the alignment of reads from several loci on the same genomic region, due to paralog collapsing during genome assembly). Finally, the maternal allele had to be present in at least one F1 read and both parental alleles had to be present in at least 75% of the F1 reads covering the SNP (both pools combined).

Finally, preliminary results revealed a problem in which some contigs showed very low probability of absence of recombination with the SDR in a given pool *i*, *P*(*f* = 0.5| ***cr***_***i***_), due to rare group-2 SNPs leading to aberrant *c*_2*i*_/*r*_2*i*_ ratios. Accurate estimate of *P*(*f =* 0.5 | ***cr***_***i***_) is critical as we use it to select contigs that may contain the SDR (see section “Localization of genomic regions that may contain the SDR”). We attribute the negative influence of “suspicious” SNPs on *P*(*f =* 0.5 | ***cr***_***i***_) to mapping or assembly errors (*c*_2*i*_ being much too high to result from sequencing errors). These errors would lead to reads from different loci aligning on the same genomic region. Hence, the apparent variation between reads would not represent allelic variation (SNPs), but another type of variation. To locate these suspicious SNPs, we computed *P*(*f* = 0.5| ***cr***_***i***_) on each individual SNP as if it constituted its own haplotype, and ignored SNPs yielding much lower posterior probability than the rest of the SNPs of each contig (see Supplementary text, Figure S2).

### Contig assignment to sex chromosomes and analysis of recombination

We used our estimate of the number of oocytes that underwent recombination between a target contig and the SDR during both crosses (*n*_rec_, equation 3) to isolate contigs that are significantly closer to the SDR than expected assuming an autosomal location. To account for the uncertainty in *n*_rec_ and approximations of our method (namely in equation 2), we contrasted the observed values of *n*_rec_ to those obtained by simulating sequencing data in the F1 pools. These simulations used the actual genomic positions of the retained informative SNPs and the identifiers of reads covering these SNPs, only changing the bases that reads carried at SNPs to reflect a given genetic distance to the SDR.

The simulations consisted in the following procedure, which we applied to every contig. First, we assigned the contig a genetic distance (in Morgans) to the SDR, which we call *d*. We then applied the following in each family independently. We designated the two homologous maternal haplotypes at the contig with letters Z and W, denoting the sex chromosome hosting these haplotypes in the mother. For each SNP that was informative in the family, the maternal allele was randomly attributed to haplotype Z or W (i.e., the SNP is attributed to type 2 or 1) with the same probability. We then applied the following to each of pool *i* of the family. We defined as *n*_*Zi*_ the number of chromosomes carrying haplotype Z in the pool of 2*n*_*i*_ chromosomes. To simulate linkage to the SDR, *n*_Z*i*_ was sampled from *Binomial*(*n*_*i*_, *d*) if the pool contained daughters or from *n*_*i*_ – *Binomial*(*n*_*i*_, *d*) if the pool contained sons. To simulate the sequencing of the maternal haplotypes, each read of maternal origin was randomly attributed to haplotype Z or W with probabilities *n*_*Zi*_/(2*n*_*i*_) and 0.5 − *n*_*Zi*_/(2*n*_*i*_), respectively. At each SNP, each read was set to carry the maternal allele if the read and the maternal allele of this SNP were both attributed to the same haplotype (Z or W). Otherwise, the read was set to carry the paternal allele. Based on these artificial reads, we inferred SNP type and computed *P*(*f* | ***cr***_***i***_) as for the real data. We repeated the procedure 1000 times for every contig, with *d* set to 0.5 to simulate autosomal contigs. If, for any contig, the value of *n*_rec_ obtained from the real data was lower than the 1/1000^th^ quantile of values obtained from simulations, we considered the contig as located on the sex chromosomes with a 1/1000 risk of false positive.

We also performed simulations to investigate a potential reduction of recombination rate near the SDR, which may evolve during sex chromosome divergence. To this end, we assessed how the cumulated length of contigs (a proxy for physical distance on chromosomes) varies according to the inferred distance to the SDR (*n*_rec_). We did so for observed data and for simulated data assuming uniform recombination rates along sex chromosomes. Uniformity in recombination rates was ensured by attributing every contig a value of *d* sampled from *Uniform*(0, 0.5). We assigned these “simulated contigs” to sex chromosomes as we did for observed data. After discarding contigs not assigned to sex chromosomes, we randomly sampled a number of simulated contigs, ensuring that their total length was as close as possible to the total length of observed contigs assigned to sex chromosomes. We repeated this sampling 1000 times to build confidence intervals of the cumulated length of contigs as a function of the inferred genetic distance to the SDR.

### Investigation of heterozygosity

The SDR and its genomic context are predicted to show increased allelic divergence compared to autosomal regions due to balancing selection combined with potentially reduced recombination. We investigated this hypothesis by measuring female heterozygosity in the two individually genotyped mothers. For each mother, we ignored SNPs of genotype quality <40, those of sequencing depth <5 or higher than the 95% quantile for the genotyped individual (only considering positions reported in the vcf file). For each contig, we counted the number of unique heterozygous positions passing these filters when considering both mothers combined. To estimate SNP density given the sequencing effort, we recorded the number of unique contig positions belonging to the aforementioned range of sequencing depths for each contig, again combining both mothers. Sequencing depth was computed with samtools, excluding duplicate reads, secondary alignments, alignments with mapping quality zero and bases with PHRED score <10, to mimic the parameters used by the SNP caller as much as possible.

We then analyzed how female heterozygosity varied according to the inferred genetic distance to the SDR (*n*_rec_). For these analyses, we opted to ignore contigs for which *n*_rec_ could not be inferred with sufficient certainty. We did so by discarding contigs for which the highest posterior probability of *f*, max(*P*(*f* | ***cr***_***i***_)), was lower than 0.5 in any pool *i*. Doing so considers that contigs with fewer informative SNPs, hence with lower female heterozygosity on average, were less likely to be assigned to sex chromosomes due to reduced statistical power. This bias might lead to a spurious correlation between female heterozygosity and assignment to sex chromosomes. Selecting contigs for which that data allowed estimating *n*_rec_ with a certain level of confidence should mitigate this bias.

### Localization of genomic regions that may contain the SDR

Our crossing scheme enabled a two-level selection of genomic regions that may contain the SDR. The first level consisted in selecting contigs that showed little evidence for recombination with the SDR during our crosses. We based our selection on the posterior probability of absence of recombination, which we multiplied across pools for each contig as follows

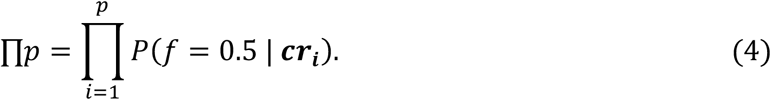

We selected contigs for which ∏*p* exceeded 0.5. We also considered selecting contigs that were more likely to have undergone 0 recombination with the SDR, i.e. contigs for which *n*_rec_ was closer to zero than to one (*n*_rec_ <0.5). This criterion was fulfilled by all the contigs retained by the former except three (109 vs. 112) and did not retain any additional contig. We opted for the more inclusive criterion.

We then used our ability to reconstruct the parental haplotypes of the selected contigs for the WXa and ZM lines to assess whether a target genomic region recombined with the SDR during the divergence of the two lines (Figure 2). This task relied on SNPs whose alleles can be assigned to parental W or Z chromosomes. These were the informative SNPs that we retained, whose alleles were assigned to maternal chromosomes via inference of SNP type, or SNPs that were homozygous in mothers (Figure 2). For each SNP that was not informative in at least one family, we imposed a minimal genotype quality of 40 and a maximum sequencing depth equaling the 95% quantile of this variable, for each parent of the family (or families). We then discarded SNPs for which the rarer allele was carried by only one parental chromosome among the four parents, as such SNPs cannot inform on recombination. We refer to the set of retained SNPs as “selected SNPs”. Recombination between a selected SNP and the SDR was inferred if these two loci constituted four different haplotypes in the parents, considering the SDR as a biallelic locus with Z and W alleles. We refer to selected SNPs that recombined with the SDR as “recombinant SNPs” (Figure 2).

**Figure 2.**
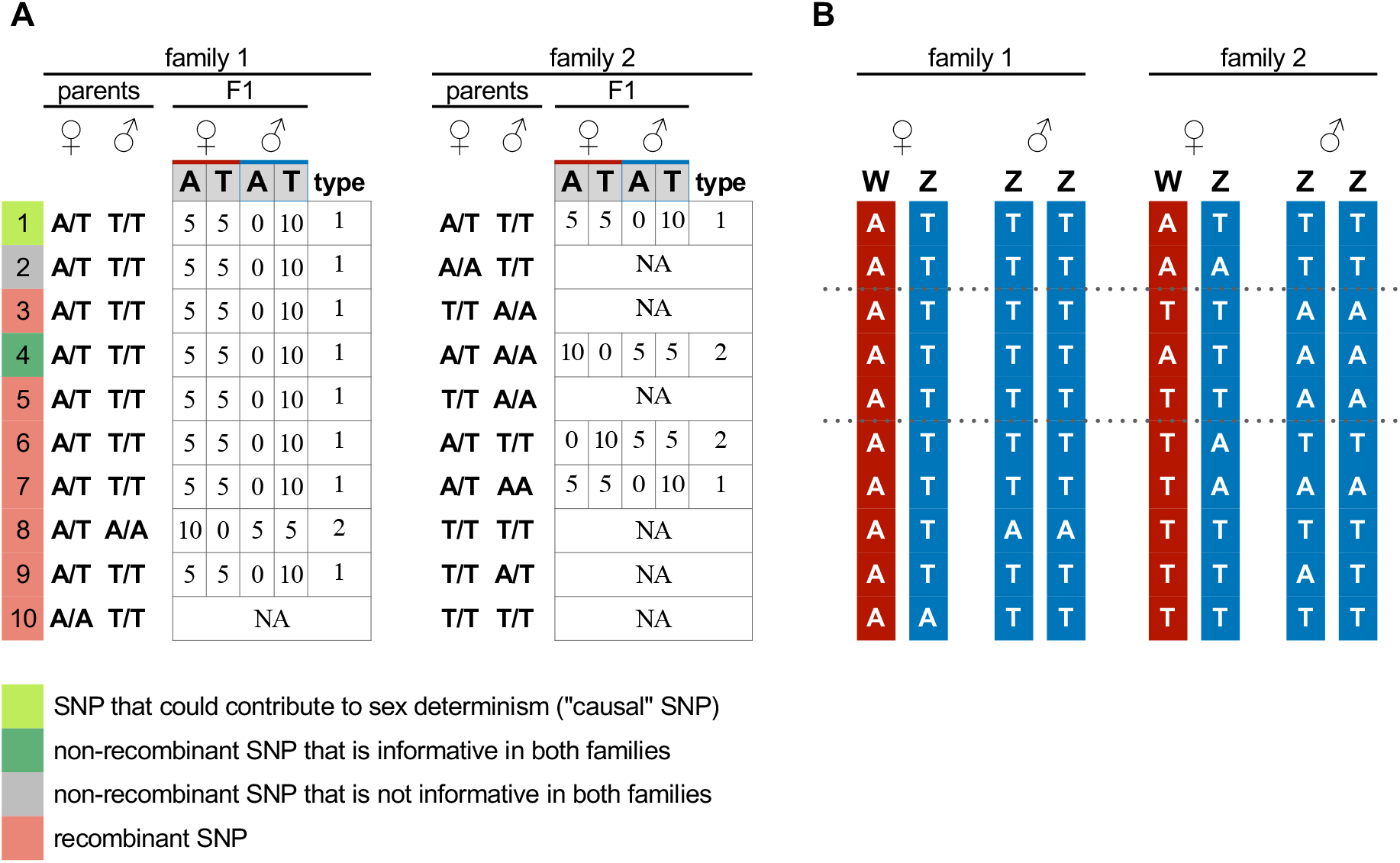
Ten hypothetical SNPs at a locus that has not recombined with the SDR during crosses involving two families. Letters A/T indicate DNA bases (alleles) at the SNPs. A) parental genotypes and data from F1 pools. Numbers (0, 5, 10) in the F1 tables indicate the number of reads that carry each allele and are only recorded for informative SNPs (otherwise, “NA” is noted). SNP type is deduced from allele frequencies in the F1 (see section “General approach”). B) Reconstructed parental haplotypes shown as vertical rods, using inferred SNP types for heterozygous mothers. Recombination must have occurred between certain SNPs and the SDR during the divergence of families, barring homoplasy in the SNPs. SNP #4 may not have recombined with the SDR, but it is flanked by two SNPs that must have. Because there is no evidence of recombination between SNP #4 and these two others (these three SNPs constitute three different haplotypes, not four), they constitute a single genomic block delineated by the doted lines.

Among selected SNPs, we looked for non-recombinant SNPs that were informative in both families (e.g., SNPs #1 and #4 on Figure 2). We reasoned that close linkage to the SDR should maintain female heterozygosity by balancing selection, hence favor this category of SNPs. Within this category, we more specifically looked for SNPs that may directly contribute to sex determination, i.e., SNPs that could compose the sex-controlling locus. Such a SNP must be heterozygous in all females and homozygous in all males, for the same bases and in all families, considering both parents and F1s. These SNPs were easily identified as being of type 1 and presenting the same maternal and paternal alleles across families (e.g., SNP #1 on Figure 2). Hereafter, we call them “causal SNPs”.

To more reliably determine that a given SNP is causal, we estimated the probability that this SNP has not recombined with the SDR in our third family (BF). Because parental genotypes were missing, we assumed that the candidate SNP was informative in this family, was of type 1, and presented the same maternal and paternal alleles as the WXa and ZM families. We computed *P*(*f* = 0.5| ***cr***_***i***_) in each BF pool *i* for each selected SNP as if it constituted its own haplotype. Not fulfilling the assumptions or having recombined with the SDR would result in a low posterior probability. We then multiplied the probabilities obtained from the two BF pools and discarded the SNP as causal if this product was <1%. This threshold was chosen after considering that 98.9% of the informative SNPs carried by the studied contigs in the WXa and ZM families had a value exceeding 1% at this variable. We therefore considered the 1% threshold as rather permissive in the selection of causal SNPs.

Beyond SNPs, we aimed at defining larger regions that may or may not have recombined with the SDR. To do so, we delineated contig regions (hereafter called “blocks”) within which no SNP showed evidence for recombination with any other, by applying the aforementioned four-haplotype criterion on every possible pair of selected SNPs (see Supplementary text for details). Note that a causal SNP and a recombinant SNP cannot be in the same block as the former behaves exactly as the ZW SDR.

We ignored every block containing a single selected SNP, unless this SNP was causal, as we considered such a short block as possibly resulting from a genotyping error rather than from recombination. As blocks were initially delineated by SNP coordinates, we extended block boundaries up to contig edges, or up to midpoints between consecutive blocks, as appropriate. We then considered any block harboring at least two recombinant SNPs or at least 50% of recombinants among selected SNPs as having recombined with the SDR.

### Search of W-specific sequences

Z-linked and W-linked alleles may not show detectable homology due to excessive molecular divergence. This possibility implies that the most divergent parts of the W and Z chromosomes, potentially including the SDR, constitute different contigs in the reference genome assembly, with some contig(s) containing W-specific regions and other contigs (or another contig) containing Z-specific regions. Such regions would not present informative SNPs since W-derived and Z-derived reads would not map on the same locations. However, the sequencing depth of a W-specific region should be close to zero for male-derived sequencing reads and it should be higher for female-derived reads. We used this criterion to locate such regions.

Sequencing depth of all six F1 pools was measured with samtools, using the same mapping quality threshold as that used for the SNP analysis. For each pool, sequencing depth was averaged over 2-kb sliding windows that moved by 500 bp, leading to a 1500-bp overlap between successive windows. To standardize results given differences in sequencing effort, we multiplied depths by the highest mean sequencing depth over the six pools (averaged over all windows) and divided them by the mean sequencing depth of the pool under consideration. We then computed the “Chromosome Quotient” (CQ) (HALL *et al.* 2013) by dividing the sequencing depth obtained from sons by that obtained from daughters, for each window in each family.

## Results

### Sex chromosomes constitute at least 43 Mbp of the *A. vulgare* genome

For the WXa and ZM families combined, more than 5.1 million individual positions of the genome were homozygous in fathers and heterozygous in mothers, constituting potentially informative SNPs. The ~3.7 million SNPs that passed our filters were carried by 40640 contigs constituting ~96.7% of the genome assembly length (1.72 Gbp). Among these, 30875 contigs (~82.2% of the genome assembly length) carried informative SNPs in both families.

Sequence data from the F1 pools at retained SNPs were used to compute *n*_rec_ (equation 3), the estimated number of oocytes that have recombined with the SDR in our crosses. Detailed results for each contig are provided in File S1. Figure 3 shows the distribution of the percentage of recombinant oocytes (*n*_rec_/40×100), which is the inferred distance to the SDR in cM. The distribution obtained from real data is similar to that obtained from simulations assuming autosomal contigs but is slightly shifted to the right, possibly due to errors that we did not simulate (Supplementary text). Despite this slight right shift, a tail is visible to the left, denoting contigs located closer to the SDR than expected for autosomes. In particular, 1004 contigs, representing the pale blue distribution in Figure 3 and totaling ~43.6 Mbp, present significantly lower distances to the SDR than expected from autosomal contigs and were assigned to sex chromosomes.

**Figure 3.**
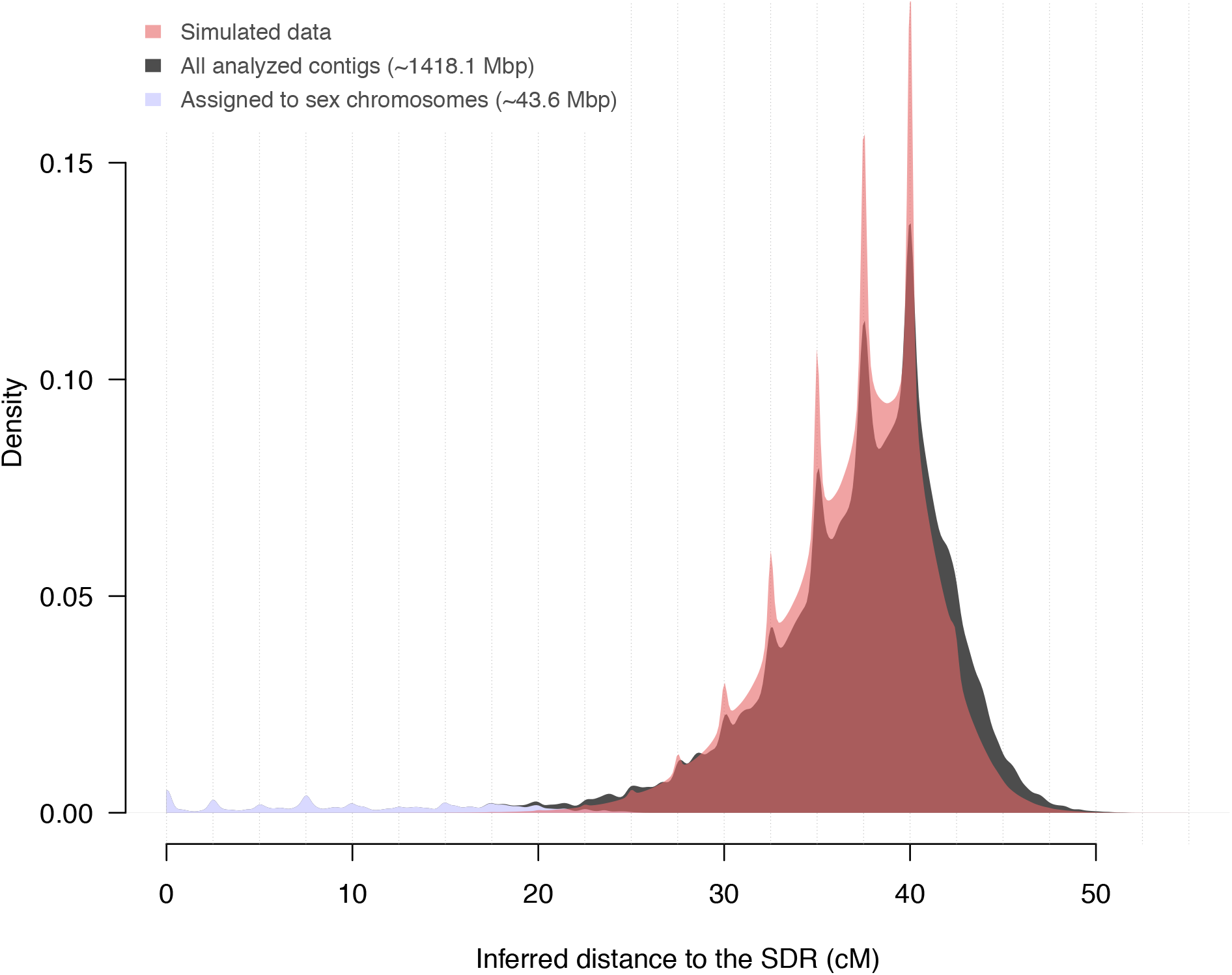
Distributions of the inferred genetic distance of *A. vulgare* contigs to the SDR for real data and for simulated data assuming that all contigs locate on autosomes. Genetic distances are inferred from simulated or observed genetic data from 40 F1 siblings belonging to two families. Vertical dotted lines represent genetic distances corresponding to integer numbers of recombination events during the crosses. Distributions only consider contigs for which both families present informative SNPs (33406 contigs). Contigs whose inferred distance to the SDR was lower than distances yielded by all simulations were assigned to sex chromosomes and constitute the blue area (see text). The distribution modes are lower than 50 cM, despite this value being the expectancy for autosomal contigs, because haplotypes cannot be properly reconstructed for contigs that are distant to the SDR (Supplementary text).

Considering that these results implicate ~82.2% of the genome assembly (contigs showing informative SNPs in both families), we extrapolate that ~53 Mbp of contigs (43.6/0.822) locate on sex chromosomes. These contigs should include about one thousandth of the autosomal contigs (false positives) at our significance level. However, false negatives are likely to be more frequent than 1/1000 according to additional simulations we performed to evaluate our method (Supplementary text, Figure S3). Therefore, the estimated length of *A. vulgare* sex chromosomes can be considered as conservative.

### The sex determining region of *A. vulgare* is located within less than 1 Mb

The sex chromosomes of *A. vulgare* appear to show relatively uniform crossing over rates. The cumulated length of contigs assigned to sex chromosomes indeed increases regularly with the inferred genetic distance to the SDR and remains within the 99% confidence intervals obtained from simulations assuming uniform crossing over rates (Figure 4). There is consequently no evidence for a reduction in crossing over rates near the SDR, in which case the cumulated length of contigs would have been higher than expected at short genetic distances.

**Figure 4.**
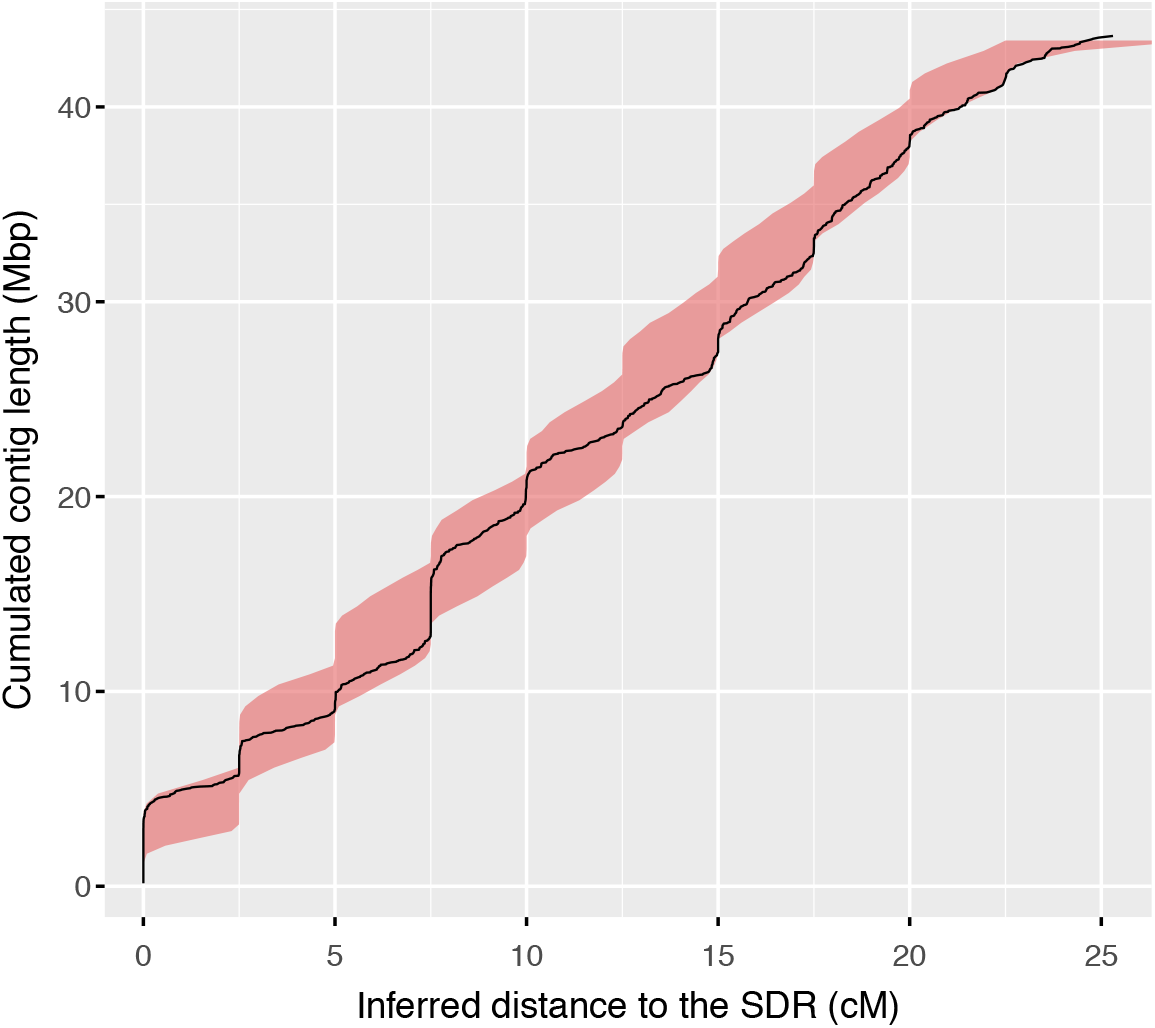
Cumulated length of 896 *A. vulgare* contigs that locate below or at a given genetic distance from the sex determining region (SDR). Genetic distances are inferred from genetic data from 40 F1 siblings belonging to two families. The black curve represents observed data and the reddish area the 99% confidence intervals obtained from simulated data assuming uniform crossing over rates along sex chromosomes.

Among the 1004 contigs assigned to sex chromosomes, 112 collectively accounting for ~5.1 Mbp (Figure 5) present little evidence of recombination with the SDR during our crosses (∏*p* > 0.5, equation 4). However, our SNP-based analysis leveraging the use of the two *A. vulgare* lines (WXa and ZM) indicated that most of these contigs have recombined with the SDR at some point after the divergence of the WXa and ZM chromosomes. Indeed, the 112 selected contigs were largely composed of genomic blocks harboring recombinant SNPs. Overall, few genomic regions (~895 kbp total) did not show evidence of recombination with the SDR. We detected only ten SNPs that could directly contribute to sex determination (i.e. potential causal SNPs for which all males appear to be homozygous and females heterozygous for the same bases) in all three families (120 SNPs if we ignore the BF family). The 10 SNPs correspond to five genomic blocks located on as many different contigs, and total ~64 kbp. None of these SNPs locates in an exon.

**Figure 5.**
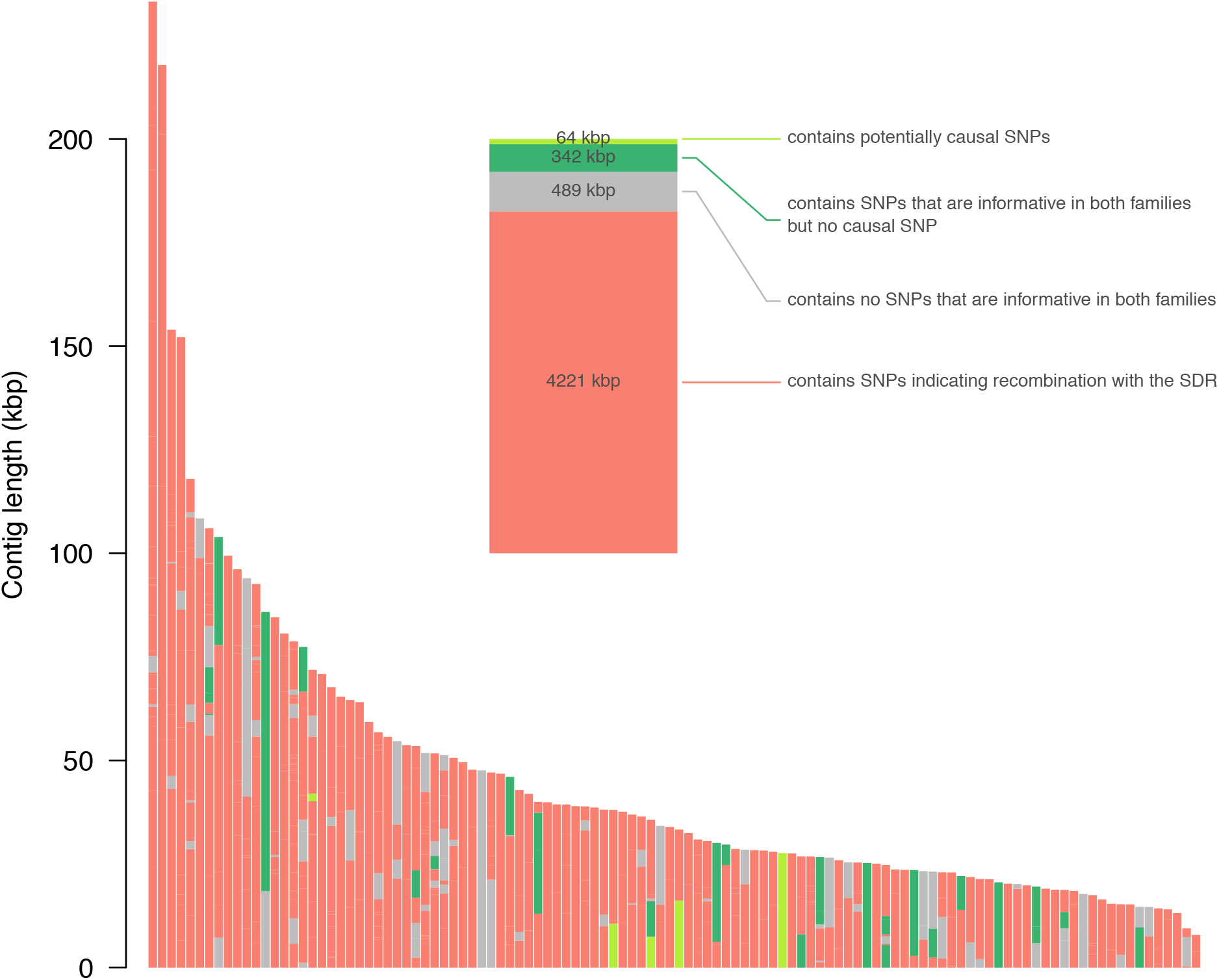
The 112 *A. vulgare* contigs that are inferred not to have recombined with the SDR in 40 F1s from two crosses. Each sectored vertical bar of the larger plot represents a contig. Contigs are ranked according to their length. Sectors within bars represent the genomic blocks compositing contigs (see section “Localization of genomic regions that may contain the SDR”). Colors represent the SNPs that genomic blocks carry, using the same color codes as in Figure 2 and in the inset. The inset shows the total lengths of different categories of genomic blocks according to the SNPs they carry. Blocks belonging to first three categories (from the top) contain no more than one recombinant SNP and less than 50% of recombinant SNPs.

The SDR had a modest impact on the molecular divergence of the Z and W chromosomes. Indeed, female heterozygosity did not significantly decrease with the inferred genetic distance to the SDR (one-sided Spearman’s rank sum test *S* = 74540329, *p* = 0.3416) (Figure 6). Yet, female heterozygosity increased at the closest distance to the SDR. Its median was indeed significantly higher for the 112 contigs that showed little evidence of recombination with the SDR than the other contigs assigned to the sex chromosomes (~10.32 SNPs/kbp vs ~9.01 SNPs/kbp, one sided Mann-Whitney’s *U* = 41839, *p* ≈ 0.005). Despite this difference, the median female heterozygosity of the contigs assigned to sex chromosomes did not differ from that of the other contigs (two-sided *U* = 5598974, *p* = 0.3253). It should be kept in mind that these comparisons do not include the whole genome as we discarded contigs for which *f* could not be inferred with a posterior probability of at least 0.5 (see section “Investigation of heterozygosity”).

**Figure 6.**
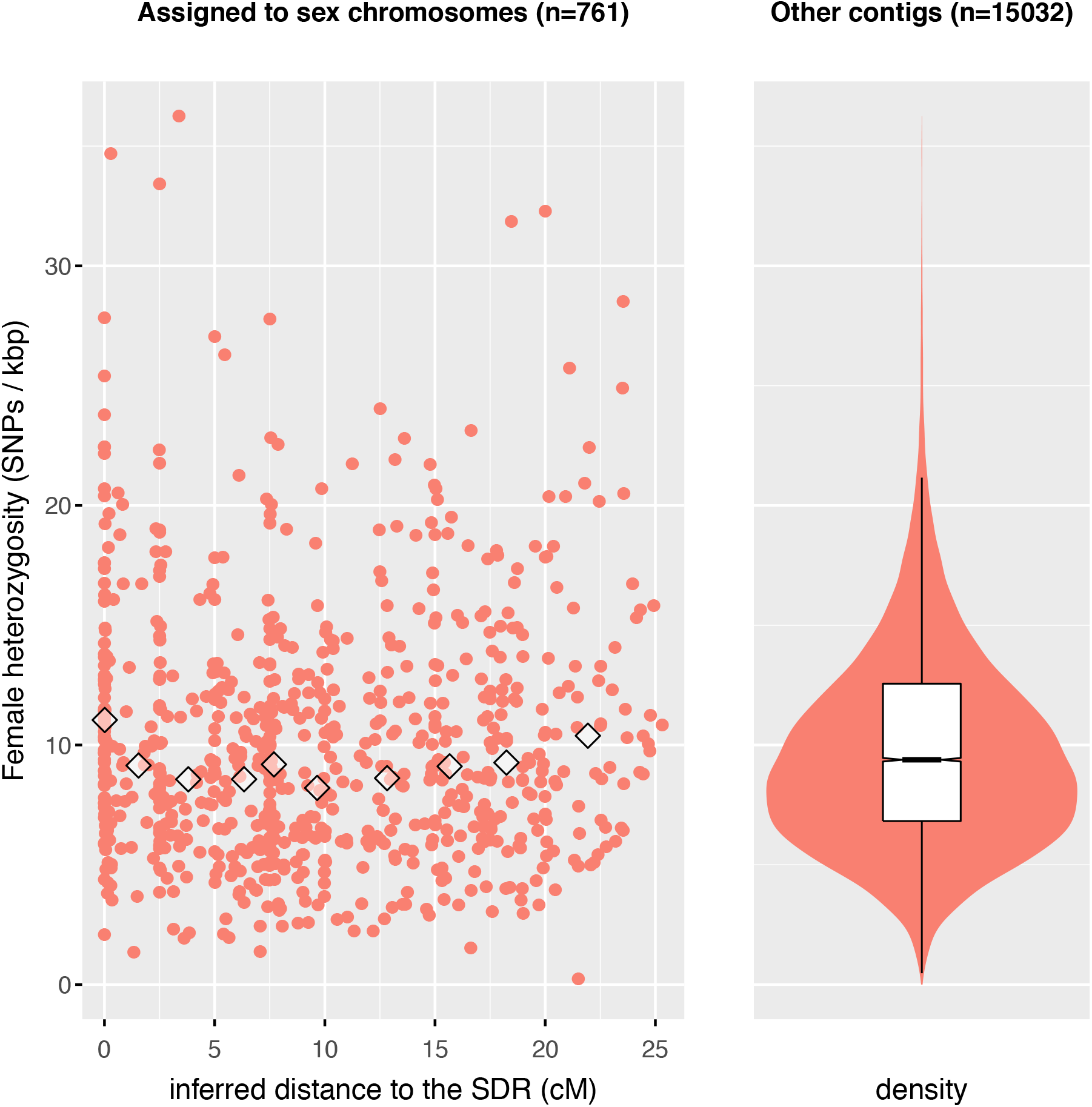
Female heterozygosity as a function of the inferred genetic distance to the SDR for contigs assigned to sex chromosomes (left-hand plot) and its distribution for other contigs (right-hand plot). The diamonds on the left-hand plot represent medians computed for ten classes of genetic distance. Classes are delimited by the deciles and therefore comprise ~70 contigs each. For the density computation (right-hand plot) contigs were weights by the lengths of regions with sufficient sequencing depth to measure heterozygosity (see methods).

### W-specific genomic regions are rare

The ratios of sequencing depth obtained from sons to that obtained from daughters (CQ scores) presented distributions that are typical of autosomes for all three families (Figure S4), showing little evidence for sex chromosomes heteromorphy. Only seven contigs contained W-specific sequences, as defined by genomic windows with CQ <0.3 and female sequencing depth >5 in all three families. These windows added up to ~92 kbp. Informative SNPs present in these seven contigs indicated that six have recombined with the SDR during the crosses, with a minimum *n*_rec_ of ~3.5. The one exception was contig 20397 (Figure 7). Remarkably, it is also the contig that possesses the most potentially causal SNPs (*n* = 4) in a single block spanning the whole contig (Figure 5). Information about the two genes in contig 20397 (Figure 7) is provided in Table 2. A DNA sequence similarity search using blastn (CAMACHO *et al.* 2009) with default settings showed that the exons of these genes were similar to exons of other annotated genes of the *A. vulgare* genome. These genes therefore present paralogs. The sequence identity of the most similar copy of each gene (Table 2) was higher than the identity measured with the most similar annotated gene of *A. nasatum* (BECKING *et al.* 2019), which was inferred to have diverged from the *A. vulgare* lineage ~25 million years ago (BECKING *et al.* 2017). This result suggests relatively recent duplications of the two genes.

**Table 2.** Annotated genes in contig 20397 containing genomic windows of low chromosome quotient in the three *A. vulgare* lines.

| Name | Number<br>of exons | Exon<br>length | Description | Number of<br>paralogs<sup>a</sup> | Highest identity with<br>paralogs<sup>b</sup> |
| --- | --- | --- | --- | --- | --- |
| AV_16253 | 2 | 204 | Hypothetical protein | 11 | 97.0% |
| AV_16254 | 2 | 228 | Hypothetical protein | 2 | 98.7% |

<sup>a</sup> genes that harbor a region of at least 100 bp having a blastn e-value <10<sup>-4</sup> with the coding sequence of the gene listed in the first column. <sup>b</sup> Identity considers the alignments reported by blastn, not the whole gene lengths.

**Figure 7.**
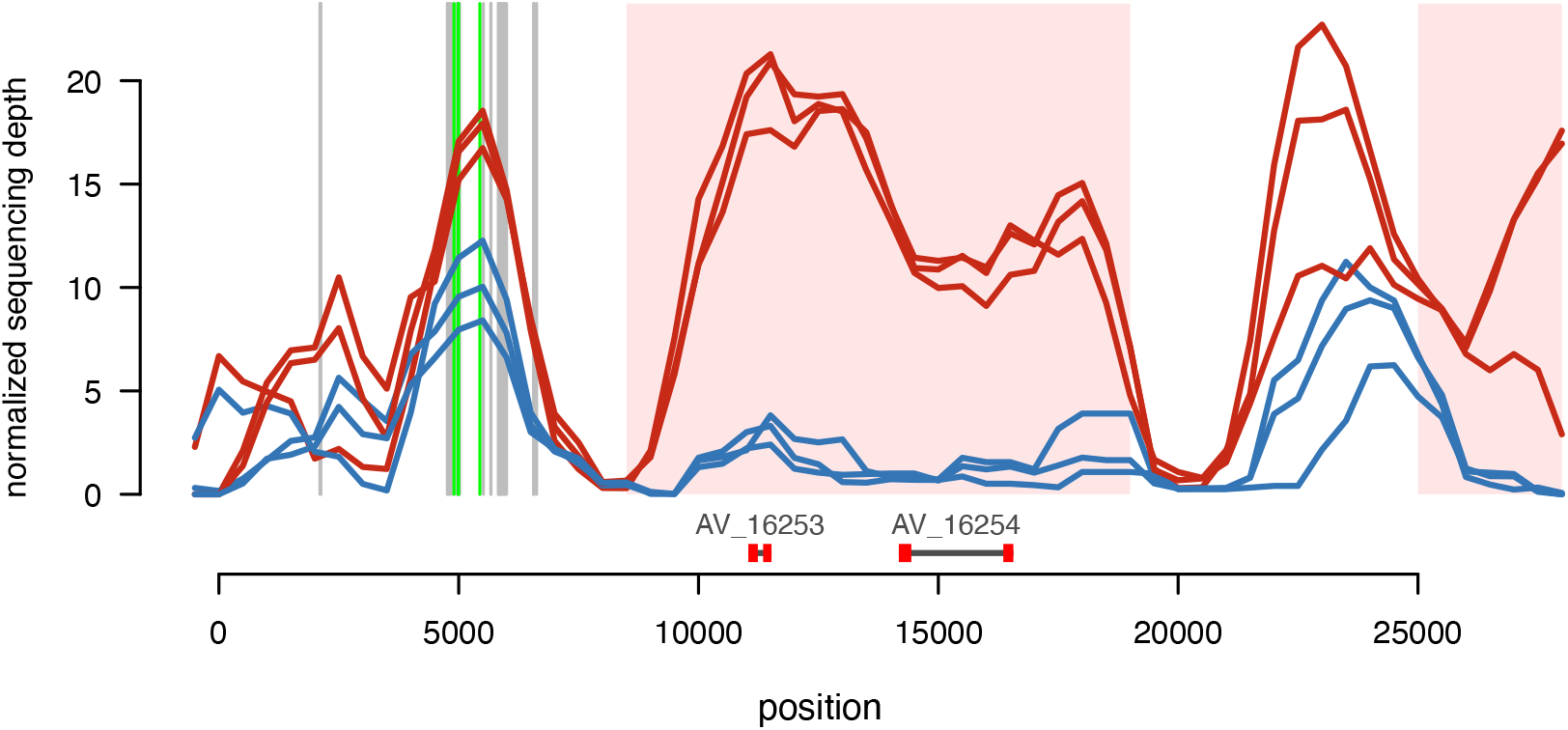
Normalized female (red curves) and male (blue curves) sequencing depths on contig 20397, presenting low chromosome quotient (CQ), based on sequenced DNA from the progeny of three *A. vulgare* families, constituting 6 pools. Regions of CQ < 0.3 and of female sequencing depth ≥ 5 in all families are represented as light pink areas. Vertical segments represent informative SNPs. Grey segments represent SNPs that are variable in a single family. Bright green segments represent SNPs that are compatible with the control of sex (“causal SNPs”, see text). Annotated genes are represented by horizontal dark grey segments under gene identifiers. Exons are shown as thick red segments.

## Discussion

### Benefits of our method

Segregation analysis methods aimed at locating sex-associated regions generally rely on individual genotyping by partial genome sequencing, using RAD tags or similar techniques as cost-saving measures (PALMER *et al.* 2019). Pool-seq appears to be less utilized for this task, especially compared to its popularity in population genomics analyses using field samples (KOFLER *et al.* 2011). However, when obtaining a genetic map is not essential, whole-genome pool-seq can be advantageous, not only because of its ease of use and applicability to any species, but mostly because of the millions of SNPs this approach can yield. In comparison, partial genome sequencing using RAD tags yields tens of thousands of reliable SNPs at most. If this number does not largely exceed the number of contigs in the genome assembly, as might be the case for a non-model organism, a high risk of false negative may affect the selection of contigs that can contain the SDR. RNA sequencing (as used in (MUYLE *et al.* 2016)) would also suffer this problem if exon density is low (~1.4% of the genome assembly in the case of *A. vulgare*), added to the fact that only a subset of genes is expressed during an experiment. The millions of SNPs yielded by whole-genome sequencing is leveraged by our approach through the combination of all the informative SNPs of a contig into haplotypes, whose frequencies are more precisely inferred than allele frequencies based on single SNPs. A high SNP density is also particularly useful to pinpoint the SDR within contigs that did not recombine with this locus during the crosses, thanks to a multi-family setup (Figure 2, Figure 5). Here, the resolution corresponds to the typical distance between SNPs that inform on recombination with the SDR and would be quite limited if partial genome sequencing were employed. In fact, our approach can be used to locate a genomic region controlling any qualitative trait that depends on a single locus. Doing so would simply require treating the phenotype that associates with the heterozygous genotype as the ZW females of our study.

We emphasize that the number of read pairs covering all the informative SNPs of a contig (variables *r*_1*i*_ and *r*_2*i*_), not sequencing depth per se, determines the certainty of the estimated allele frequencies. Contigs with more informative SNPs, hence longer contigs for a given SNP density, thus require lower average sequencing depth for a similar degree of certainty in estimated haplotype frequencies. While larger F1 pools should yield improved imprecision, the size of a pool, *n*, must ensure that 1/*n* is at least twice as high as the sequencing error rate, to clearly differentiate rare alleles from errors. Cost considerations aside, a larger number of families (hence of pools) can circumvent this limitation and also increase the ability to detect past recombination with the SDR during the divergence of studied lineages, hence to exclude genomic regions as containing the SDR.

The precision yielded by whole-genome pool-seq extends to the location of regions harboring sequences that are specific to the rarer sex chromosomes (here, the W) via the CQ analysis. The use of sliding windows, which would not be permitted under partial genome sequencing, allows low-CQ regions to be shorter than contigs. In this case, the statistical analysis of nearby SNPs provides a useful complement to the CQ scores. Indeed, the absence of reads having a particular DNA sequence from one sex (CQ=0) may not imply the complete absence of this sequence from the genome analyzed individual(s), due to the odds of DNA sequencing. Also, it is unclear whether a low CQ value indicates low genetic similarity between sex chromosomes at the focal genomic window (allowing a small portion of reads from a chromosome to map on the sequence of the other) or the rare occurrence of an allele in the sex where this allele is supposed to be absent. The analysis of nearby SNPs reduces these uncertainties by providing a probabilistic assessment of the association between genetic variants and sexes, which may extend to low-CQ regions of the same contig. This combination of approaches allowed us to outline contig 20397 among the seven contigs showing low-CQ windows in *A. vulgare*.

### Extremely low divergence of *A. vulgare* sex chromosomes

Seven contigs of the *A. vulgare* genome assembly contain ~92 kbp of potential W-specific sequences, and only contig 20397 was inferred to not have recombined with the SDR during our crosses. Our previous study (CHEBBI *et al.* 2019) outlined 27 contigs containing W-specific sequences, only two of which (including contig 20397) presented low-CQ windows in the present study. This difference can be explained by the fact that CHEBBI *et al.* (2019) investigated a single family, reducing the probability of recombination with the SDR, and computed CQ at the scale of whole contigs rather than genomic windows. Indeed, 14 of the 27 contigs outlined by CHEBBI *et al.* (2019) are among those we assigned to sex chromosomes (considering that four of the 27 contigs did not have informative SNPs and could not be assigned). Due to partial linkage to the SDR, these 14 contigs would have harbored genetic differences that associated with the sex of individuals studied in CHEBBI *et al.* (2019), but this association did not hold in the three lines we analyzed, except for contig 20397. The low CQ scores reported by CHEBBI *et al.* (2019) at these contigs therefore do not amount to sex chromosome divergence, but to polymorphism in these chromosomes. The other nine contigs that we did not assign to sex chromosomes may be more distant to the SDR. Their median inferred genetic distance to the SDR of ~31.5 cM is still significantly lower than the median of other contigs not assigned to sex chromosomes (39.3 cM) (two-sided Mann & Whitney’s *U* = 64752, *p* ≈ 8.5×10^−4^). Thus, some of the nine contigs may belong to sex chromosomes despite not having passed the assignment test.

The scarcity of W-specific sequences in the *A. vulgare* genome extends to Z-specific sequences. Indeed, the CQ distributions obtained from the three studied lines (Figure S4) show no particular bump near 2, which is the expected CQ value for Z-specific regions. We did not specifically study regions with high CQ, because elevated CQ on short genomic windows is subject to high sampling variance, hence to false positives/negatives, as opposed to low CQ which involves low sequencing depth, hence low variance. At any rate, these results demonstrate the very low divergence of the *A. vulgare* sex chromosomes. In our previous study (CHEBBI *et al.* 2019), the inference of low divergence was not definitive as the size of sex chromosomes was undetermined. Here we show that the sex chromosomes have a minimal size of 53 Mbp, that is, 83% the average size of *A. vulgare* chromosomes (~64 Mbp) based on genome size and number of chromosomes. Even though our estimate of sex chromosome length is conservative, it is orders of magnitude that of W-specific sequences.

The reduced divergence of *A. vulgare* sex chromosomes is consistent with their recombination rates. Only ~5.1 Mbp of contigs would not have recombined with the SDR during our crosses, and most have recombined with the SDR since the divergence of the ZM and WXa lines (Figure 5). The fact that sex chromosomes did not show evidence for uneven crossover rates (Figure 4) and are not distinguishable from autosomes in terms of female heterozygosity (Figure 6) suggests comparable levels of recombination between chromosome types. The higher density of heterozygous SNPs in contigs locating the closest to the SDR (Figure 6) could just be the byproduct of balancing selection, increasing coalescent times of SDR-linked alleles over a relatively short genomic region. Hence, we do not consider this observation as conclusive evidence for a reduction of crossing over rate around the SDR.

The apparent absence of recombination reduction in *A. vulgare* sex chromosomes could reflect their recent origin, considering the apparent rapid turnover of sex chromosomes in terrestrial isopods (BECKING *et al.* 2017). An evolutionary scenario for this renewal involves feminizing bacterial endosymbionts of the genus *Wolbachia* (RIGAUD *et al.* 1997; CORDAUX *et al.* 2011). *Wolbachia* endosymbionts that infect terrestrial isopods can improve their maternal transmission by feminizing their carriers, as commonly observed in *A. vulgare* populations (JUCHAULT *et al.* 1993; VERNE *et al.* 2012; VALETTE *et al.* 2013). Theoretical models and field surveys on *A. vulgare* indicate that invasion of a ZW population by feminizing *Wolbachia* leads to the loss of W chromosomes, as feminized ZZ individuals take the role of mothers (reviewed in CORDAUX AND GILBERT (2017)). Sex then becomes entirely determined by the presence of *Wolbachia* and is generally biased towards females, reflecting the prevalence of the feminizing bacteria. New sex chromosomes may emerge via the selection of masculinizing nuclear genes that may reestablish an even sex ratio (CAUBET *et al.* 2000; BECKING *et al.* 2019) or through horizontal gene transfer of feminizing *Wolbachia* genes into host genomes, producing new W-type chromosomes (LECLERCQ *et al.* 2016; CORDAUX AND GILBERT 2017).

With these considerations in mind, we cannot strictly exclude that the ZW chromosomes of *A. vulgare* are old, as female heterogamety prevails among Armadillidiidae and related families (BECKING *et al.* 2017). The *A. vulgare* sex chromosomes may have been maintained as homomorphic for a long period through sustained recombination, similarly to what is observed in palaognaths (XU *et al.* 2019), European tree frogs (STÖCK *et al.* 2011) or guppies (DAROLTI *et al.* 2020). Characterizing the SDRs of ZW species that are closely related to *A. vulgare* should allow evaluating the homology, hence the age, of sex chromosomes among these lineages.

### The sex determining region of *A. vulgare*

Barring false negatives, the sex-controlling locus of *A. vulgare* should lie within the ~0.9 Mbp of genomic blocks that did not show evidence for recombination with the SDR (Figure 5). As these blocks span several contigs that also harbor recombinant SNPs, the total length of the SDR, as defined by the non-recombining chromosomal region surrounding the master sex-determining gene(s), is likely to be much shorter than 0.9 Mbp. Where the SDR locates in these 0.9 Mbp cannot yet be determined, but we expect this locus to harbor Z-specific and W-specific alleles that have been maintained for a long time by balancing selection. We therefore do not favor the 489 kpb of genomic regions harboring no SNP whose alleles could be associated to the W and Z chromosomes in both *A. vulgare* lines. On the opposite, the 64 kbp of genomic regions containing causal SNPs are of greater interest as they contain sex-associated variation in all three investigated families. Among the candidate genomic regions, contig 20397 clearly emerges. This contig harbors the region with the highest number of causal SNPs, no SNP indicating past recombination with the SDR and a low-CQ region encompassing two annotated genes. The low CQ on these genes indicates the presence of large indels and/or the accumulation of smaller mutations that make male reads unable to align on the reference sequence (i.e., the allele on the W chromosome). As the two annotated genes of contig 20397 appear to be female-specific, their expression may be required for development into a female. Interestingly, these genes present highly similar paralogs. Evolution of master sex-determination genes by duplication of existing genes has been reported in diverse taxa, such as medaka fish (MATSUDA *et al.* 2002; NANDA *et al.* 2002) and clawed frog (YOSHIMOTO *et al.* 2008). Unfortunately, the available functional annotation for the two genes on contig 20397 (Table 2) precludes any speculation about possible mechanisms of action. Moreover, the coding sequences of the two female-specific genes in *A. vulgare* present at most six substitutions with their closest paralog, preventing any reliable inference about natural selection acting on their evolutionary branch. We therefore cannot exclude that these genes are redundant copies that happen to be linked to a nearby feminizing allele, and whose absence in males has no consequence. We also keep in mind that a ZW sex determining system may not necessarily involve feminizing gene transcripts or proteins, and that sex determination in *A. vulgare* could depend on expression levels of Z-specific genes, as in the chicken (SMITH *et al.* 2009). Investigating these hypotheses requires searching for Z-specific sequences in the *A. vulgare* genome. While CQ scores resulting from comparisons of ZZ and ZW individuals may not be reliable for this task, as previously discussed, CQ scores obtained by comparing laboratory produced WW individuals (JUCHAULT AND LEGRAND 1972) and ZZ or ZW individuals should be close to zero for Z-specific sequences, similarly to the CQ scores we used to locate W-specific sequences.

To conclude, the extremely low molecular divergence of the *A. vulgare* sex chromosomes and their apparently uniform recombination rates allowed us to pinpoint a limited set of regions that could contain the SDR and to identify two potential feminizing genes. Their strict association with the female sex in additional *A. vulgare* lines and their expression levels during sex differentiation will be the focus of future research. An improved reference genome assembly would also lessen the risk that the sex-determining locus was missed due to insufficient sequencing coverage and/or number of informative SNPs, which mostly affect short contigs. By mapping our sequence data on a more contiguous (chromosome-scale) genome assembly and applying our approach on long genomic windows, the SDR of *A. vulgare* should be characterized in its entirety.

## Data availability

The custom code used to perform the data analysis is given in File S2. The genetic data generated in this study are deposited to Genbank under accession numbers [to be deposited].

## Acknowledgements

We thank Clément Gilbert for discussions at an early stage of the project, Bouziane Moumen for providing support for our computing infrastructure and the genotoul bioinformatics platform Toulouse Midi-Pyrénées (bioinfo.genotoul.fr/) for providing computing and storage resource. This work was funded by European Research Council Starting grant 260729 (EndoSexDet) and Agence Nationale de la Recherche grant ANR-15-CE32-0006-01 (CytoSexDet) to R.C., the 2015–2020 State-Region Planning Contracts (CPER) and European Regional Development Fund (FEDER), and intramural funds from the Centre National de la Recherche Scientifique and the University of Poitiers.

## Supplementary text

### How SNP type inference conditions the estimates

SNP type designates the chromosome (Z or W) carrying the allele that is specific to the mother (see section “General approach”). SNP type is inferred by comparing the frequency of this allele between daughters and sons, which permits the reconstruction of maternal haplotypes whose frequencies are estimated in the F1 pools. This inference relies on linkage between a locus and the SDR. In fact, the notion of SNP type is irrelevant with respect to autosomes.

To assess how SNP type inference affects the estimated number of recombination events with the SDR, *n*_rec_, we performed simulations as described in section “Contig assignment to sex chromosomes and analysis of recombination” (main text). In these simulations, every contig was attributed a genetic distance to the SDR, *d*, covering every value of the set {0…20/40}. *n*_rec_ was estimated as in equation 3 (main text) and contrasted with the number of recombinants that we simulated (which is the sum of *n*_*Zi*_ in daughters and (*n*_*i*_ − *n*_*Zi*_) in sons). We also recorded the proportion of SNPs whose type was properly inferred. As a basis of comparison, we performed equivalent simulations in which SNP type was not inferred but taken directly from the simulated SNP type.

Results are shown in Figure S1. When the simulated SNP type is used to compute *n*_rec_ (Figure S1A), *n*_rec_ provides a good estimate of the actual number of recombinants at all genetic distances to the SDR. When SNP type is inferred (Figure S1B), *n*_rec_ tends to underestimate the true number of recombinants. The underestimate coincides with errors in SNP type inference (Figure S1C). Expectedly, inference of SNP type does not perform better than a random guess at 20 recombinants (corresponding to 50 cM to the SDR) considering that types 1 and 2 are equifrequent in the simulations.

Why *n*_rec_ is underestimated comes from the fact that the “focal haplotype” regroups the maternal alleles of group-1 SNPs, which, given how SNP groups are defined, are more frequent (among reads) in the sex under consideration than in the other. If the two maternal haplotypes are roughly equifrequent among sexes of an F1 progeny, as expected for an autosomal locus, our method would create a chimeric focal haplotype gathering the maternal alleles that happen to prevail among reads, and whose inferred frequency will *de facto* be higher than that of real haplotypes. The focal haplotype appearing more frequent in a sex, the number of recombination events with the SDR is underestimated.

### Exclusion of SNPs that may result from errors

For a contig containing many informative SNPs in a pool *i*, a single SNP should have little influence on *P*(*f =* 0.5 | ***cr***_***i***_), hereafter called *p*_05_ and representing the posterior probability of absence of recombination with the SDR. In some scenario however, only one or few SNPs can almost nullify *p*_05_, which would otherwise be close to 1. This scenario involves an SDR-linked contig for which all SNPs are of group 1 the pool. Due to errors, some “suspicious” SNPs may occur and may be attributed to group 2 if the maternal alleles appear more frequent than in the other pool of the same family. The strong influence that these few group-2 SNPs have on *p*_05_ arises from the fact that a high value of *c*_2*i*_ (leading to *c*_2*i*_ /*r*_2*i*_ largely exceeding the sequencing error rate ε) is incompatible with *f* = 0.5. On the other hand, *f* = 0.5 involves a high variance in *c*_1*i*_ (this variance equals 0.25*r*_1*i*_ under a binomial distribution). As a result, a single (false) group-2 SNP can easily nullify *p*_05_ even if hundreds of group-1 SNPs support *f* = 0.5.

This risk prompted us to find and exclude suspicious SNPs resulting from errors. To do so, we computed *p*_05_ on each individual SNP as if it constituted its own haplotype (that is, the read counts *c*_1*i*_, *r*_1*i*_, *c*_2*i*_ and *r*_2*i*_ only consider the reads covering this SNP, such that *r*_1*i*_ or *r*_2*i*_ is zero). We reasoned an apparent SNP yielding a value of *p*_05_ that is much lower than the majority of SNPs of the same contig should result from an error. We excluded these SNPs with the following iterative procedure, which was applied separately to each contig in each pool. We exclude any SNP for which −ln(*p*_05_) was higher than the mean of this variable added to four times its standard deviation, both computed over on all non-previously excluded SNPs of the contig. We repeat this until no further SNP was excluded. Figure S2 illustrates the approach.

### Interpretation of the distributions of *n*_*rec*_ for real and simulated data

The distributions of the inferred distance of contigs to the SDR (Figure 3, main text) show two features that we discuss here.

First, the distributions are lower than the expectancy of 50 cM for autosomes. This is due to the fact that *n*_rec_ underestimates the true number of recombination events at large distances to the SDR (see section “How SNP type inference conditions the estimates”).

Second, the distribution of the observed genetic distances appears shifted to the right compared to simulations. We suggest that this shift reflects uncontrolled errors. In the absence of error, *c*_1*i*_ /*r*_1*i*_ and *c*_2*i*_ /*r*_2*i*_ should negatively correlate, since their respective expectancies are *f* and 0.5−*f*. This is not so if the maternal alleles of a locus represent errors (genotyping error in parents, sequencing errors in F1s etc.) where we expect the ratios to positively correlate, as they represent the error rates. For instance, if the apparent maternal alleles result from errors, *c*_1*i*_ /*r*_1*i*_ and *c*_2*i*_ /*r*_2*i*_ ratios may both be low. Similar *c*_1*i*_ /*r*_1*i*_ and *c*_2*i*_ /*r*_2*i*_ ratios for a contig result in an inferred value of *f* closer to 0.25, hence increase the inferred distance from the SDR.

### Evaluating the power of assignment tests to sex chromosomes

To evaluate our method of contig assignment to sex chromosomes, we simulated data as described in section “Contig assignment to sex chromosomes and analysis of recombination”. We gave every contig a value of *d* sampled from the set {0…20 / 40} to cover a range of distances to the SDR. As we did for observed data, a contig was assigned to sex chromosomes if the obtained *n*_rec_ was lower than a low-order quantile of *n*_rec_ obtained from simulations that used *d* = 0.5. The proportion of contigs assigned to sex chromosomes among those located at <50 cM to the SDR represents the power of the assignment test, i.e., the sensitivity of our approach.

To evaluate how the sensitivity of the method depended on its specificity, the order of the aforementioned quantile, which is the risk of false positive or significance level, was allowed to vary in the set {0.001, 0.005, 0.01, 0.05}. Note that for contig assignment using observed data, the 1/1000^th^ quantile was used.

We also investigated how the reconstruction of maternal haplotypes, via inference of SNP type, affected the sensitivity of our approach. To this end, we performed a similar set of simulations in which SNP type was not inferred but directly deduced from the chromosome carrying the maternal allele, which is set during the simulation (see main text). In this case, reconstructed haplotypes correspond to simulated haplotypes.

Figure S3A shows how the ability to assign contigs to sex chromosomes depends on the distance from the SDR, the significance level and on whether SNP type was inferred. At the 0.001 significance level and when SNP type is inferred (as it was the case for observed data), sensitivity falls under 60% at 20 cM to the SDR.

Figure S3B shows the proportion of contigs assigned to sex chromosomes among those locating at or below a certain distance to the SDR, assuming uniform crossing over rates along sex chromosomes. For instance, ~86% of contigs locating at ≤20 cM to the SDR are assigned to sex chromosomes at the 0.001 risk.

The reduction of sensitivity due to lack of knowledge on SNP type (represented by the distance between plain and dotted lines on the Y axis) is higher at intermediate genetic distances (Figure S3A). This is due to the fact that SNP types are properly inferred for contigs that locate close to the SDR (see section “How SNP type inference conditions the estimates”). At high genetic distances, knowing the true SNP types provides little benefit as the high number of crossing overs make distant contigs similar to autosomal contigs.

When it comes to the number of contigs properly assigned among those locating at a certain distance or closer to the SDR (Figure S3B), not knowing SNP type incurs a loss of sensitivity of eight percentage points at most (~72% vs. ~64% sensitivity at 32.5 cM).

### Delineation of genomic blocks based on parental genotypes

The “genomic blocks” (see section “Localization of genomic regions that may contain the SDR”) are contig regions within which no SNP shows evidence for recombination with another. Each block is considered as a unit when it comes to the possible location of the SDR. To delineate blocks, we proceeded as follows. We first generate every possible pairs of SNPs in a contig, using those we refer to as “selected SNPs”. We then check whether a pair of SNP constitutes four different haplotypes among parents, in which case, recombination must have occurred between the SNPs. To generate these two-SNP haplotypes, we use knowledge of the chromosomes (either the W or Z) carrying the alleles of informative SNPs for heterozygous mothers (this knowledge is permitted by inference of SNP type, see main text). We do not reconstruct the haplotypes of a father that is heterozygous at both SNPs, as we do not know the chromosomes carrying the SNP alleles.

We then scan the SNP pairs to delineate the longest continuous regions containing no pair of recombinant SNPs. These regions often overlap (Figure S5A). We removed overlaps by iteratively shortening the shortest region that overlapped with any other (Figure S5B).

## Supplementary figures

**Figure S1.**
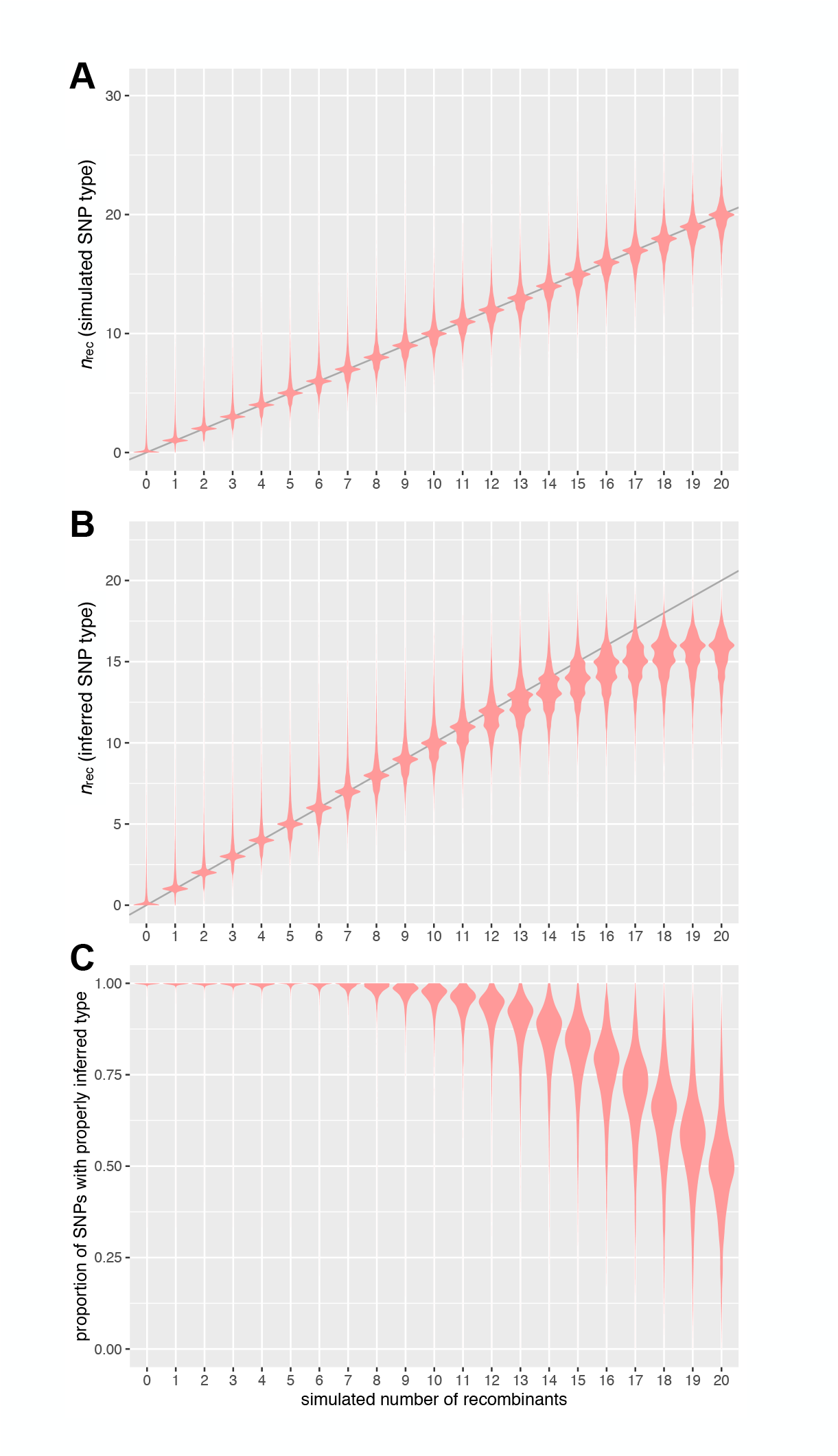
Results of simulations assessing the accuracy of the estimate of the number of recombinants with the SDR. The data points constituting the distributions are contigs possessing informative SNPs in both families. In A), the number of recombinants between the SDR and a contig was estimated without inference of SNP type, but using knowledge of the simulated SNP type. In B), the estimates rely on the inference of SNP type. Panel C) shows the proportion of SNPs whose type was properly inferred, combining both families.

**Figure S2.**
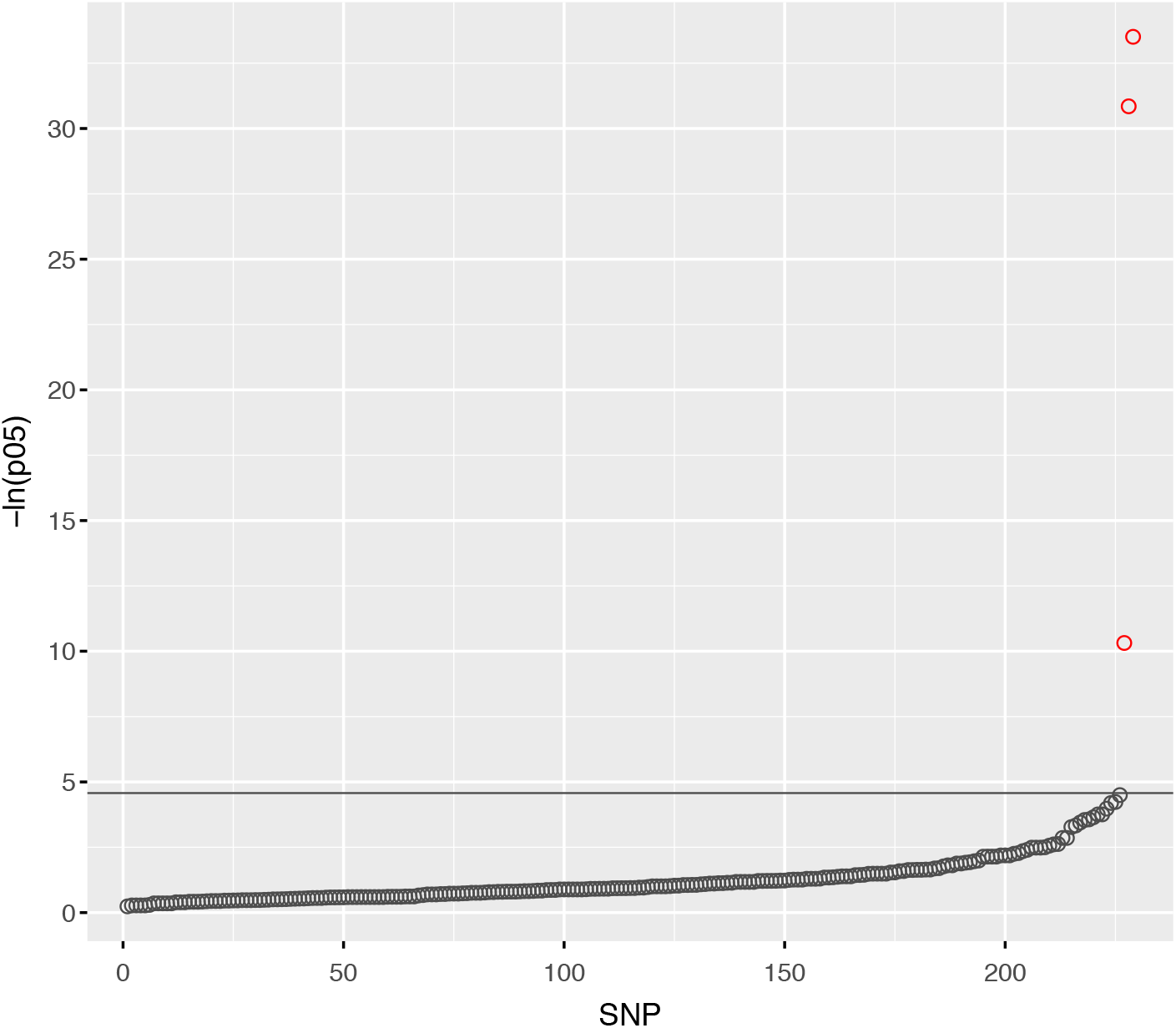
–natural logarithm of *p*_05_ (the probability that *f* = 0.5) for individuals SNPs of contig 28198 in daughters from the WXA family. SNPs are ordered by increasing −ln(*p*_05_). Group-1 SNPs are shown as dark grey points, group-2 SNPs as red points. The horizontal line represents the threshold above which SNPs are excluded. Excluding the three group-2 SNPs result in *p*_05_ ≈ 0.99 for the whole contig. Not excluding these SNPs results in *p*_05_ ≈ 0.

**Figure S3.**
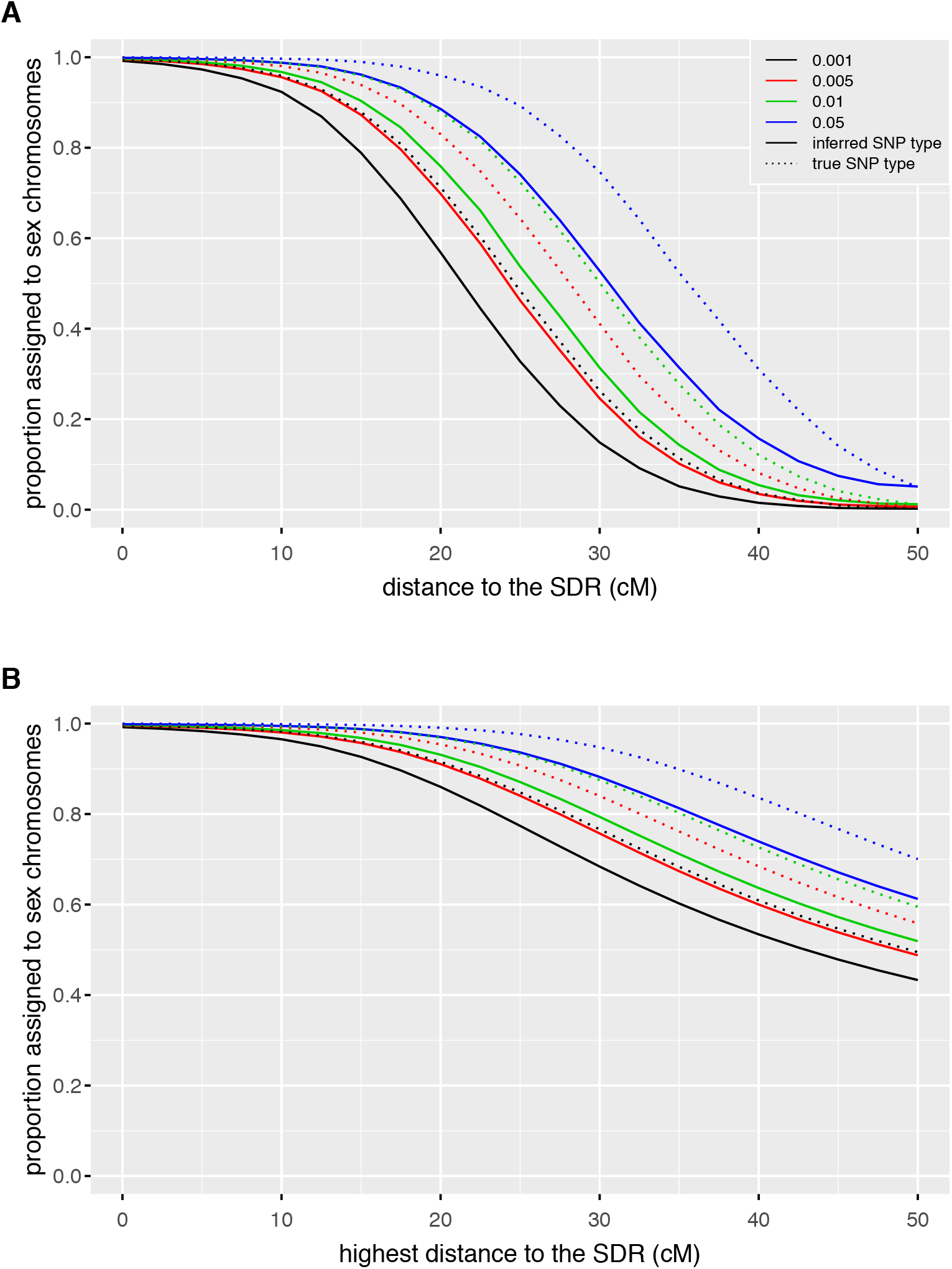
Power of the assignment of contigs to sex chromosomes. A) Proportion of contigs assigned to sex chromosomes as a function of the simulated genetic distance between contigs and the sex-determining region (parameter *d*, here expressed in cM). B) Proportion of contigs assigned to sex chromosomes among contigs whose simulated genetic distance equals or is lower than the value of the X axis, assuming that genetic distance to the SDR varies linearly with the number of contigs. In both panels, colors represent the different significance levels used, and line styles indicate whether SNP types were inferred of directly taken from values set during simulations (see supplementary text).

**Figure S4.**
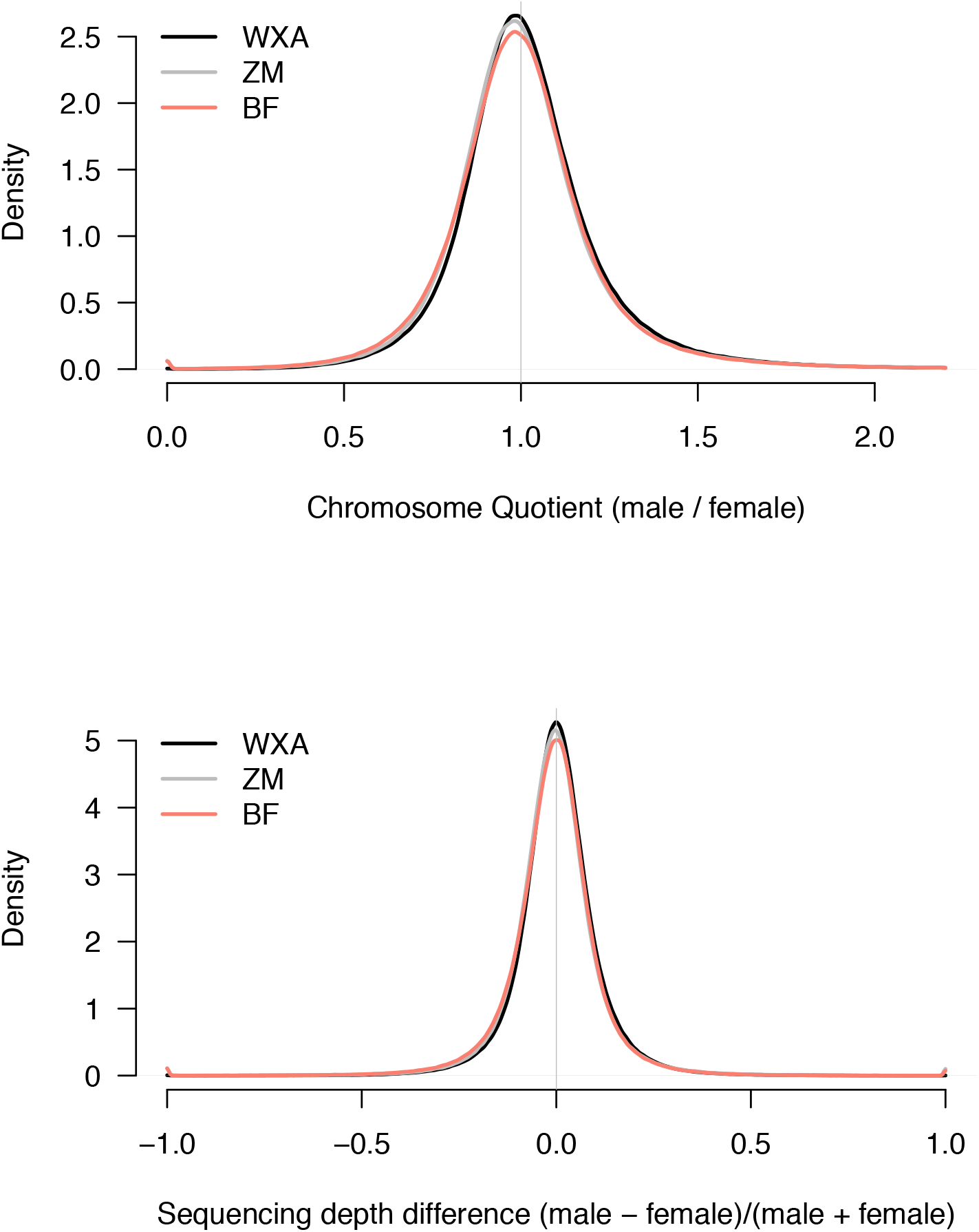
Comparison of normalized sequencing depths between sons and daughters within 3 families of *A. vulgare*, whose names are shown in the plot legends. Sequencing depths are averages computed for 2kpb genomic windows sliding by 500pb. Windows of mean sequencing depth < 3 (summed within a family) are excluded to reduce noise. The Chromosome Quotient (top) presumably has a mode < 1 due to the right skewness of the variable, which varies from zero to infinity. This does not indicate that normalized sequencing depths is typically lower in males. The variable used in the bottom plot does not have this issue as it ranges from −1 to 1. Its mode is ~0 and extreme values are very rare, indicating that most genomic windows have similar sequences between sexes.

**Figure S5.**
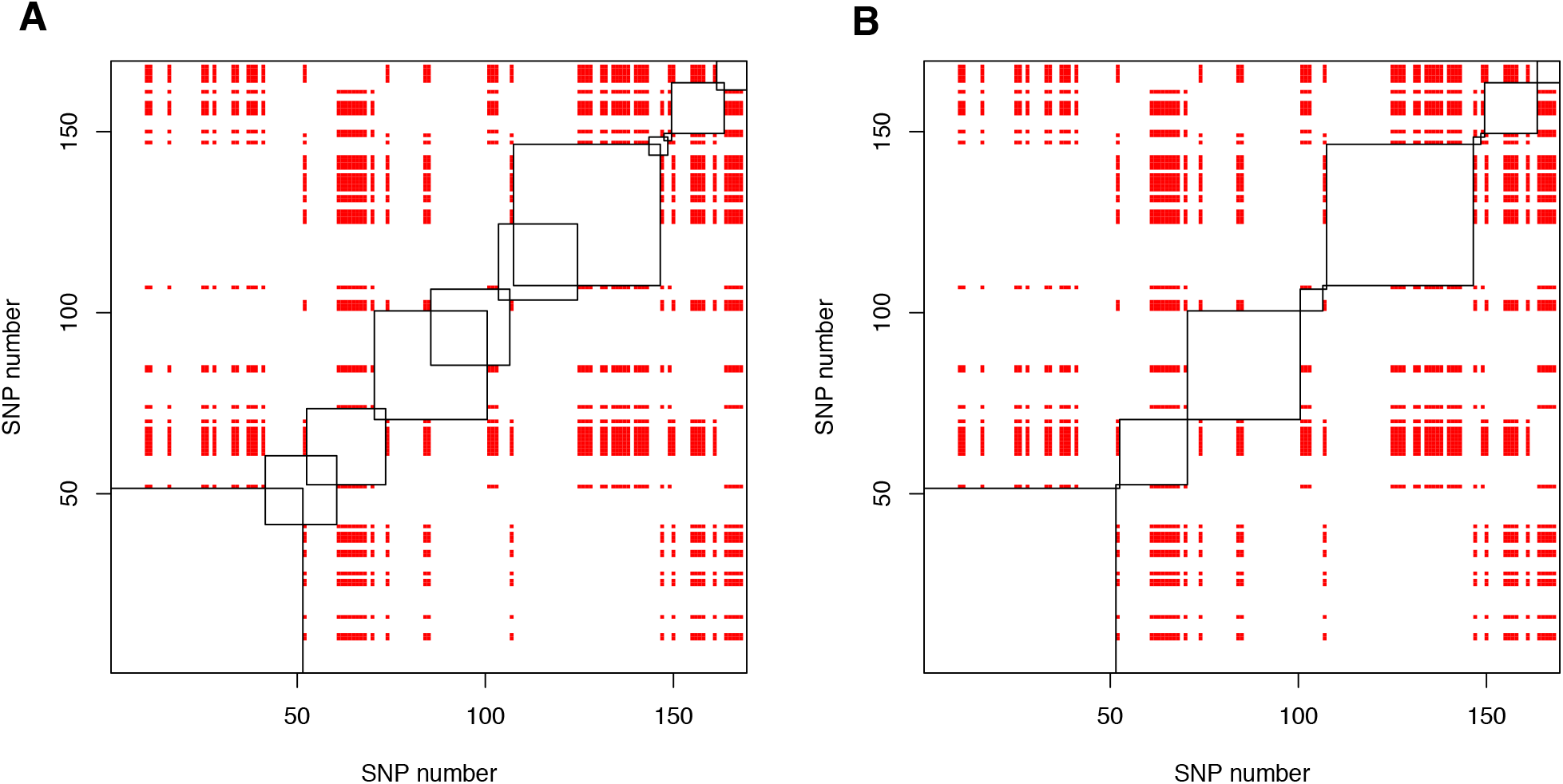
Delineation of genomic blocks in contig 42878, which contains 169 SNPs used to infer recombination with the SDR. In each image, a row/column represents a SNP according to its relative position in the contig. Each “pixel” is therefore a pair of SNPs. A red pixel denotes evidence for recombination between the SNPs of a pair. The ascending diagonal can be viewed as the 169 SNPs. Squares represents genomic blocks (see text). In A), the initial blocks are overlapping. In B), the smallest blocks were iteratively shortened to avoid overlaps. The three shortest blocks are removed afterwards as they contain only one selected SNP each.

## Supplementary files

**File S1.** This excel file contains information about each contig analyzed. For each family, the posteriori probability of the focal haplotype frequency *f* being 0.5 is given, together with the posterior probability of the most likely haplotype frequency, and the read counts. Other columns include the contig name, its length, its inferred genetic distance to the SDR (*n*_rec_), whether it is assigned to sex chromosomes, and the female heterozygosity.

**File S2.** Compressed archive of the custom code used in this study.

